# Similarity corpus on microbial transcriptional regulation

**DOI:** 10.1101/219014

**Authors:** Lithgow-Serrano Oscar, Gama-Castro Socorro, Ishida-Gutiérrez Cecilia, Mejía-Almonte Citlali, Tierrafría Víctor, Martínez-Luna Sara, Santos-Zavaleta Alberto, Velázquez-Ramírez David, Collado-Vides Julio

## Abstract

The ability to express the same meaning in different ways is a well known property of natural language. This amazing property is the source of major difficulties in natural language processing. Given the constant increase in published literature, its curation and information extraction would strongly benefit by efficient automatic processes, for which, corpora of sentences evaluated by experts is a valuable resource. Given our interest in applying such approaches to the benefit of curation of the biomedical literature, specifically about gene regulation in microbial organisms, we decided to build a corpus with graded textual similarity evaluated by curators, and designed specifically oriented to our purposes. Based on the predefined statistical power of future analyses, we defined features of the design including sampling, selection criteria, balance, and size among others. A non-fully crossed-design was performed for each pair of sentences by 3 evaluators from 7 different groups, adapting the SEMEVAL scale to our goals in four successive iterative sessions with a clear improvement in the consensuated guidelines and inter-rater-reliability results. Alternatives for the corpus evaluation are widely discussed. To the best of our knowledge this is the first similarity corpus in this domain of knowledge. We have initiated its incorporation in our research towards high throughput curation strategies based in natural language processing.

## 1 Introduction

Expressing the same approximate meaning with different wordings is a phenomenon widely present in the every day use of natural language. It shows the richness and polymorphic power of natural language, but it also exhibits the complexity implied in understanding the conveyed meaning. Due to these characteristics, paraphrase identification is necessary in many Natural Language Processing (NLP) tasks, such as information retrieval, machine translation, plagiarism detection, among others. Although strictly a “paraphrasis” refers to a rewording stating the same meaning, a binary decision, frequently, a graded paraphrasing is needed. This graded paraphrasing is often called Semantic Textual Similarity (STS).

Textual similarity depends on particular text features, domain relations and on the applied perspective, therefore it has to be “defined” according to the context. This context specification presuppose the delineation of the *kind, of textual similarity* desired, e.g., assigning grades of importance to the syntactic parallelism, to the ontological closeness, to the statistical representations likeness, etc.

It is not a simple endeavor to explicitly state these grades of importance. Its difficulty relies on the fact that it is very complicated to envisage all possible language’s feature variations to express the same idea, and so to have a broad perspective and identify which feature or relations are important. It is here that a paraphrase corpus is a very useful instrument because it implicitly enclose those nuances.

There are several paraphrase corpora available, both in general and specific domains. However, as stated before these corpora are very sensitive to the aimed task and to the targeted domain. Hence, when a task or domain are very specific and the available corpora do not fit. an ad-hoc corpus has to be built. This is the case for the biomedical curation of the literature about the regulation of transcription initiation in bacteria, a very specific domain.

RegulonDB^1^ ([GCSSZ^+^16]) is a manually curated standard resource, an organized and computable database, about the regulation of gene expression in E. coli K-12. It aims at integrating in a single repository all the scattered information about genetic regulation in this microorganism including elements about transcriptional regulation, such as: promoters, transcription units (TUs), transcription factors (TFs), effectors that affect TFs, active and inactive conformations of TFs, TFs binding sites (TFBSs), regulatory interactions (RIs) of TFs with their target genes/TUs, terminators, riboswitches, small RNAs and their target genes. RegulonDB plays an important role in scientific research with citations in more than 800 scientific publications within the gene regulation field.

As an ongoing effort to enrich the already curated information and to improve the curation process, we are developing Natural Language Processing (NLP) tools some of which relay on STS. Due to the very specific nature of our domain we built an ad-hoc graded paraphrase corpus to be used as training and evaluation source of truth for our NLP tools.

Within the next sections we first describe the methodology followed to build our corpus, then we analyze it quantitatively and finally we briefly mention the immediate foreseen uses.

## 2 Related work and Motivation

Semantic Textual Similarity (STS) aims at measuring the degree of semantic equivalence between two fragments of texts. As so, it tries to unveil the meaning conveyed by a textual expression and compare it with the meaning conveyed by another one. The comparison’s result is a graded similarity score that ranges from an exact semantic match to a completely independent meaning, passing through a continuous scale of graded semantic parallelism. This scale intuitively captures the notion that a pair of texts can share different aspects of meaning at different levels ([ACD^+^13]), i.e. it could differ in just some minor details, could share a common topic and important details, or just share the domain and context, etc. Another characteristic of STS is that it treats similarity between two texts as bijective putting it apart from textual entailment where the relation is directed and can not be assumed in the other way around.

Many NLP tasks, such as machine translation, question answering, summarization, and information extraction, are beneficed from this quantifiable graded bidirectional notion of textual similarity. Building this kind of corpus is difficult and expensive that is why there are not as many corpora of this kind as expected given their utility.

In recent years the most notorious effort on the STS task and their correspondent corpus construction has been tackled by the *Semantic Evaluation workshop* (SEMEVAL) ([ACD^+^13]). The *SEMEVAL corpus* consists of 15,000 sentence pairs coming from different sources. *Microsoft Research Paraphrase* (MSRP) and PASCAL VOC ([EVW^+^10]) corpora among them. It was annotated through crowd sourcing on a scale from 5 (identical) to 0 (completely unrelated).

Another corpus that is useful for STS is the *User Language Paraphrase Corpus* (ULPC) ([MM11]). It was built by asking students to rephrase target sentences. As a result 1998 sentence pairs were annotated with ratings ranging from 1 to 6 for 10 paraphrasing dimensions; entailment and lexical, syntactic and semantic similarity are among those dimensions.

The SIMILAR corpus ([RLMB12]) is a qualitative assessment of 700 pairs of sentences from the MSRP corpus, and besides providing word-to-word semantic similarity annotations, it also supplies a qualitative similarity relationship identical, related, context, close, world-knowledge, and none between each pair of sentences.

Not in the scope of graded similarity but instead in the binary paraphrase one, there are important corpora such as the *Microsoft Research Paraphrase* (MSRP) ([DB05]). It is one of the first major public paraphrase corpora comprising 5801 news sentence pairs of which 67% were judged “semantically equivalent” by two human judges. In the Q&A field. *The Question Paraphrase Corpus* ([BG08]) was built by collecting, from *WikiAnswers,* 7434 sentences formed by 1000 different questions and their paraphrases.

All these corpora target general domains and were sourced mainly from news making very difficult to fit them into a such an specific topic like ours, the *bacterial transcriptional regulation.* Closer to our domain is the BIOSSES corpus ([SÖÖ17]). It is formed by 100 pairs of sentences from the biomedical domain which were rated following the guidelines of the STS SEMEVAL’s task. The candidate sentences were collected by taking those that cited the same reference article from a set of 20 reference articles each of which had another 12-20 citing articles. Articles were taken from the *Biomedical Summarization Ti-ack Tra/ining Dataset* from the *Text Analysis Conference.*

Due to the extension of the biomedical domain and the small size of this corpus, the nuances of our subject of study are not captured by it. For this reason we decided to build our own corpus of naturally-occurring non-handcrafted sentence pairs within the subject of *regulation of gene expression in E. coli K-12* literature. The semantic similarity grade of each pair was evaluated by human experts in the bacterial gene regulation domain.

## 3 Materials and Methods

### 3.1 Corpus design

“A corpus is a collection of pieces of language text in electronic form, selected according to external criteria to represent, as far as possible, a language or language variety as a source of data for linguistic research”. ([Sin04])

Before building a corpus the source textual set, the evaluation rules, the corpus size and other characteristics must be defined. This design should be, as possible, informed and principled so the resulting corpus fulfill the desired goals. The decisions taken within the axes of consideration ([Sin04]) for the corpus construction are the following.

The *sampling* policy defines where and how the candidate texts are going to be selected following 3 main criteria: the *orientation,* in this case a contrastive corpus that aimed to show the language varieties that express the same meaning (semantic similarity): the *selection criteria* that circumscribed candidates to written sentences (origin and granularity) in English (language) taken from scientific articles (type) on the topic of genetic regulation (domain) where the sentence attitude^2^ was irrelevant and a specific content was not required: finally, the *sampling* criteria consisted in a sentence pairs pre-selection through a very basic STS component and a posterior filtering process to keep the same number of exemplars for each similarity grade, i.e., a balanced candidate set.

The corpus *representativeness* and *balance* refers to the kind of features and the exemplars distribution of each of those features, hence, these characteristics determine the corpus’ usage possibilities. In this sense, sentences containing any biological element or knowledge were preferred but not limited to. It was more important that all similarity grades were represented within the corpus and preferably in equally proportions. Our main analysis axis was at the semantic similarity between a pair of sentences and not the topic represented by each sentence, neither the sentences specialization or technical level, nor the ontological specificity.

The corpus *topic* orientation impact directly in the resulting vocabulary variety and size. Whereas embracing more topics can broad the corpus possibilities of use, it can also have negative consequences in the semantic similarity measures due to the increased chances of the same term having different meanings for different topics (ambiguity). Consequently, a limited set of topics is preferred. Our intention was that the corpus be representative of the genetic regulation literature. It is worth noting that it was not limited to those sentences specifically about genetic regulation but all kind of sentences present in the correspondent literature. The corpus *homogeneity* was tackled by stripping out those sentences considered too short (less than 10 words) ([KKM]) and those sentences that were not part of the main body of the article^3^

Finally, the corpus’ *size* should be dependent on the questions that it is aimed to answer and the type of tasks where it would be applied ([PCH07], [Juc12]). However, in practice it is largely restrained according to available resources (time, money and people). Our main goals are, to train our STS system and to measure its performance. Because our STS system is based on the combination of several similarity methods, and more to come, it is difficult to estimate the required number of cases that would fulfill significant training source for the reason that it depends of each type of metric. For example, one of the most demanding methods on training data are neural networks, whose complexity can be expressed based on the number of parameters (*P*) and it is common practice to have at least *P*^2^ training cases. This would results in thousands of training cases, which is out of our reach. Thus we focused in the second goal, to measure the STS system performance. We plan to measure the Pearson’s correlation between the computed system similarity and that generated by human-experts (corpus). Accordingly to [Coh92], considering a medium size effect (r==0.30), a significance level of 0.05 and a power of 80%, 85 samples would be enough. However, [MG14] and [Pen] suggested a minimum sample size of 120 cases in order to cover not only the Pearson’s correlation but a regression analysis as well. With this in mind, we decided to generate a corpus of 170 sentence pairs, just above those thresholds.

Lastly, as a validity exercise, we compared our design decisions versus those taken in other corpora, for example, the Microsoft Paraphrase Corpus. Within the construction of the MSRP corpus ([DB05]), several constraints were applied to narrow the space of possible paraphrases. However, in our opinion and for our specific purpose, these guidelines limit the aspects of semantic similarity that the corpus could capture. For example: only those pairs of sentences with at least 3 words in common and within a range of Levenshtein edit distance were considered, but these are constraining similarity, at least at some degree, to a textual one; it was required that for a pair to be candidate, the length in words of the shorter sentence had to be more than 66% of the length of the longer sentence thus limiting the possibility for the corpus to represent the cross-level semantic similarity ([JPN16]), a phenomenon of sentences with different length. It also worth to note that the MSPRP has an agreed consensus that 67% of the proposed sentence pairs are paraphrases, meaning that the majority of sentences are semantically equivalent and, therefore, other grades of similarity and even non-similarity are under-represented.

#### 3.1.1 Compiling the corpus

As stated in the sampling criteria of the corpus design, the selection of candidate pairs was performed using a basic STS process that automatically assigned similarity scores consisting in continuous numbers between 0 and 1 inclusive, where 1 represented exactly semantic equivalence^4^ and 0 totally unrelated meaning. Next, the final candidate sentences were selected by a balanced stratified random sampling from those pre-rated sentence pairs.

This process was applied to two different sets: the *anaerobiosis FNR* subset formed by articles about the anaerobiosis from which 40% of the sentences were taken; and the *general* set, consisting of sentences taken by randomly sampling all RegulonDB’s articles from which the remaining 60% was selected. The former subset was built manually by an expert-curator who selected sentences considered relevant within the subject. The later subset was generated by randomly choosing two sentences from each of the 5963 processed articles. In order to focus on sentences belonging to the article’s main body, only sentences within the 30% to 70% of the article’s length were selected^5^.

### 3.2 Annotation design

Besides the corpus design, it was necessary to delineate the semantic similarity rating process. We followed a similar rating scale to the one used in SEMEVAL. An ordinal scale ranging from 0 to 4, where 0 represents a totally disconnected semantic relation between the two sentences, and 4 conveys an exact semantic match, having in the middle 3 other similarity shades, as shown in table 1.

**Table 1:** Rating scale

| Similarity tag | Description |
| --- | --- |
| 0 | The two sentences are on different topics. |
| 1 | The two sentences are not equivalent, but are on the same topic. |
| 2 | The two sentences are not equivalent, but share some details. |
| 3 | The two sentences are roughly equivalent, but some important information differs/ missing. |
| 4 | The two sentences are completely or mostly equivalent, as they mean the same thing. |

*Each sentence pair would be rated by 3 evaluators* selected by chance from 7^6^ different groups, which in turn were formed by the combination of 3 from 7 different human-experts. This resulted in a *non-fully crossed design,* i.e., different items (subjects) were annotated by different raters. Even when it is possible that only 2 annotators rated each item ([DLL^+^12]), we considered that 3 annotators bring the best balance and flexibility; on the one hand it allows to evaluate a larger number of exemplars and, on the other. 3 is the smallest number to use the median when there is no consensus and a discrete final score is desired.

Due to the fact that “what is considered semantically equivalent” is prone to be biased by personal subjective considerations, it was necessary to homogenize the annotation process among raters. To achieve this, a training period (annotation guideline refinement period) of 4 iterated sessions was scheduled, so annotators could get familiarized with the annotation guidelines and the corpora to be rated. During this training, each session consisted in evaluating a small subset of sentence pairs and at the end of each session, disagreements would be discussed and resolved, and annotation guidelines more precisely defined. This training period was considered concluded when a minimum annotator inter-agreement was achieved or the annotators qualitatively considered that they have fully understood the annotation guidelines.

#### 3.2.1 General guidelines

In order to make the annotation process less subjective some general guidelines were given to raters. These were collected from other corpus building experiments ([TMSP17]) and from our own observations, including:

- *Order,* clauses in compound sentences can be arranged in a different order without this implying a change in its meaning.
- *Missing clauses*, in the other, it do not automatically results in a zero similarity. It depends on grade of importance of the shared information.
- *Adjectives,* missing adjectives in principle do not affect similarity
- *Enumerations*, missing elements can be considered as a minor decrease in the similarity score unless enumeration convey the main sentence meaning. Reordering is considered as equivalent.
- *Abbreviations*, are considered equivalent, e.g. “vs” and “versus”
- *Hyperonyms and hyponyms*, share a grade of similarity “sugar substance” vs “honey” vs “bee honey”
- *Generalization or abstractions*, consider that two textual expressions share some grade of semantic similarity if one is a generalization or abstraction of the other, e.g. 8 vs “one digit number”

#### 3.2.2 Consensuated refinement

General guidelines were subsequently refined and enriched during the consensus sessions.

As a first approximation to clarify the rating scale in our context, it was decided to use the class of the Regulon’s objects as topic markers within the sentences. RegulonDB contains objects of the following classes: Gene, Features, Promoter, Transcription Unit (TU), Regulatory Interaction (RI), Reaction, Transcription Factors(TF), Growth Conditions (GC). Examples clarifying the scale similarity are the following:

- **[4]** Both sentences have in common the same objects and express the same meaning, i.e. they are paraphrase of each other. Example cases:

– Both sentences express the same meaning.

* *This would, mean that the IS5 element is able to provide FNR regulatory sites if inserted at appropriate positions.*
* *In any case, insertion of an IS5 element is able to increase FNR-dependent expression or to place genes under FNR control.*
- **[3]** Both sentences share the same objects and other elements of its meaning. However one of the sentences lack relevant elements or refers to the same objects, arrive to different conclusions. Example cases:

– Both sentences refers to the same Gene and share all other information, except that in one it is an activated and in the other is repressed.
– Sentences referencing to same RI but just differ in the RI’s conditions.
– Both sentences are almost paraphrase but one have more details.

* *These results confirm that the N-terminal domain of NikR is responsible for DNA recognition.*
* *In preliminary experiments, vie have also found that a subset of mutations within the DNA region protected by the N-terminal domain reduce the affinity of NikR for the operator data not shown!.*
- **[2]** Both sentences share at least one specific object and some other similarities. In this sense, contrasting conditions are considered as equivalent conditions, this is, they are taken as the same object. Example cases:

– Both sentences refer to the same TF

* *The fnr mutant was thus deficient in the anaerobic induction of fumarate reductase expression.*
* *Aerobic regulation of the sucABCD genes of Escherichia coli, which encode K-ketoglutarate dehydrogenase and succinyl coenzyme A synthetase: roles of Arc A, Fnr, and the upstream sdhCDAB promoter.*
– Contrasting conditions taken as equivalent conditions.

* *Aerobic regulation of the sucABCD genes of Escherichia coli, which encode K-ketoglutarate dehydrogenase and succinyl coenzyme A synthetase: roles of Arc A, Fnr, and the upstream sdhCDAB promoter.*
* *Transcription of the fdnGHI and narGHJI operons is induced during anaerobic-growth in the presence of nitrate.*
- **[1]** Both sentences have the same object’s class in common but the specific object is different. Due to Gene and GC objects are highly common in RegulonDB’s literature, it was decided that sharing just these classes is not a sufficient condition for sentences to be rated as **[1]**. When comparing sentence that refers to a TF with other that refers to any other object (or GC) that refers to the same process in which the TF is involved, the sentence pair has to be considered as **[1]**. Example cases:

– Both sentences refer to sequences and genes, even when neither the sequences nor the genes refered are the same.

* *The low level of 3-galactosidase expression seen in dinF1::Mu d (Ap lac) derivatives containing the lexA7J:TnS mutation but not the lex AS J (Def) mutation might then be due to a polar effect of the Tn,5 insertion in the lexA gene on transcription at the dinF locus.*
* *Sequence comparisons (18) suggest that E. coli and S. typhimurium have the same gene order in the metA-metH region, in agreement with P22 mapping studies.*
– Other example

* *To construct lADHop656 (operon fusion) and lADHpr656 (protein fusion), a 1.1-kb DNA fragment was excised from pADH8 by BglII and BstYI.*
\**The product was ligated into the BamHI site of the plasmid pRS415 (for an operon fusion) and pRS414 (for a protein fusion).*
- **[0]** Sentences do not even share object’s class. Example cases:

– Sentences share to Gene and GC class (exception of **[1]** rate) but not the same specific objects.

* *Aerobic regulation of the sucABCD genes of Escherichia coli, which encode K-ketoglutarate dehydrogenase and succinyl coenzyme A synthetase: roles of Arc A, Fnr, and the upstream sdhCDAB promoter.*
* *Carbon metabolism regulates expression of the pfl (pyruvate formatelyase) gene in Escheiichia coli.*

It was clarified that sentences do not necessarily have to contain biological content or refer to RegulonDB’s objects to be annotated and have rates above *0.* The annotation is assessing the similarity in meaning irrespective of the topic.

#### 3.2.3 Annotation process

To facilitate the annotation process, we decided to provide annotators with a spread sheet template (see figure 1). The template was designed so that all needed information would be in it and the rater did not had to switch to other files. It consist of a list of all sentence pairs that the annotator has to rate, for each pair the sentences’ ID and text are displayed. This area where the user writes the scores is organized by columns where each column represents an annotation session with date and time at the top cells. A small rating scale table is also included as reference.

**Figure 1:**
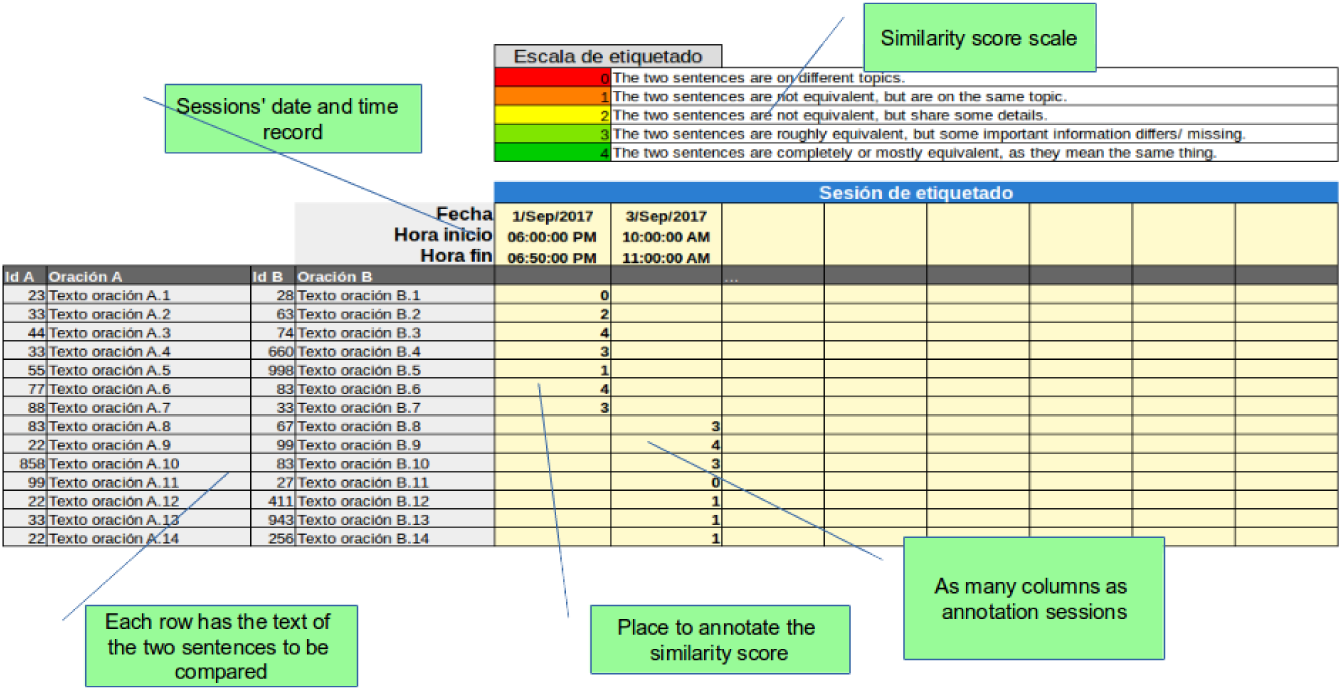
Annotation tompiate

The process consisted in: provide each annotator with a file, based on the annotation template, containing exclusively the sentence pairs that have to be evaluated by him/her; annotators had a fixed period of time of one week to rate all pairs; during that period each annotator could divide the rating task in as many sessions as desired adding date and time of each; it was indicated that sessions should be exclusive and continuous, i.e., task should not be interrupted by more than 5 minutes and annotators should not be performing other tasks in parallel.

## 4 Corpus evaluation

The recommended way to evaluate the quality of the resulting corpus is through the Inter-Rater Agreement also known as Inter-Rater Reliability (IRR) ([Hal12, Gwe02a, VBMR14, DLL^+^12, BMB08. Mch17]). IRR is a mean to measure the agreement between two or more annotators who rated an item in a nominal, ordinal, interval or ratio scale. It is based on the idea that *observed scores* (*O*) are the result of the scores that would be obtained if there were no measurement error –*true scores* (*T*)– plus the *measurement error* (*E*), i.e. *O* = *T* + *E* ([Hal12]). One possible source of the measurement errors is the measure instrument’s instability when multiple annotators are involved. IRR focuses in analyzing how much of the observed scores’ variance corresponds to variance in the true scores by removing the measurement error between annotators. Thus, the reliability coefficient represents how close are the given scores (by multiple annotators) to what would be expected if all annotators would have used the same instrument; the highest the coefficient, the better the scores reliability.

There are multiple IRR statistics and which should be used depends on the study design. Factors such as: the type of the measured variable (nominal, ordinal, etc.); if it is a fully-crossed design or not; and if what it is desired is to measure the annotator or the ratings reliability have to be considered to select the IRR statistic and/or its variant.

Our design (see section 3.2) corresponds to non fully-crossed design where an ordinal variable is measured and we are interested in measuring the ratings reliability. Having that in mind, the statistics that better accommodate are *Fleiss’ Kappa* (Fleiss) ([Fle71]). *Krippendorff’s alpha* (Kripp), *Intra Class Correlation* (ICC) ([Bar66]), *Kendall* (Kendall) ([Ken48]) and *Gwet’s AC1* (Gwet) ([Gwe02a]).

One of the most used IRR statistic is the Cohen’s Kappa analysis (k) (eq 6) ([Coh60]). It is a relation between *the proportion of units in which the annotators agreed,* 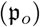 and *the proportion of units for which agreement is expected by chance* 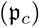, thus 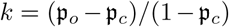. Originally this measure was intended for just two annotators who rate all items, so variants were developed in order to fit non fully-crossed study designs with more than two raters per item. The *Fleiss’ Kappa* (eq 1). is a no weighting measure that considers unordered categories and it was designed for cases when *m* evaluators are randomly sampled from a larger population of evaluators, and each item is rated by a different sample of *m* evaluators. In equation *p*_*a*_ represents the averaged extent to which raters agree for the items rate, and *p*_*ϵ*_ the proportion of assignments to the categories.

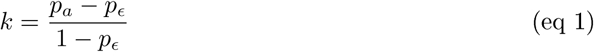

*Krippendorff’s Alpha* (eq 2) is a inter-rater reliability measure that is based on computing the disagreement. It provides advantages like being able to handle missing data, handling various sample sizes, and supports categorical, ordinal, interval or ratio measured variable metric. In eq 2, *D*_*o*_ is the observed disagreement and *D*_*ϵ*_ the disagreement one would get if rates were by chance, thus it is the ration between the observed disagreement and the expected disagreement.

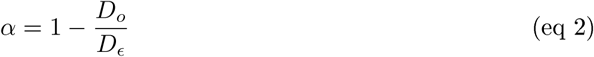

*Intra-class correlation* (eq 3) is a consistency measure that can be used to evaluate the ratings’ reliability by comparing the item’s ratings variability to the variability of all items and all ratings. It is appropriate for fully-crossed as well for non-fully crossed designs and when there are two or more evaluators. Another feature is that the disagreement’s magnitude is considered in the computation, such as in a weighted Kappa. In *var*(*β*) accounts for variability due to differences in the items, *var*(*α*) from variability due to differences in the item’s revaluations, and *var*(*ϵ*) for the variability due to differences in the rating scale used by annotators. Consistent with our study design, we selected the ICC variant as: A “one way” model, to avoid from accounting for systematic deviations among evaluators because annotators for each item are selected at random; and used the average as the unit of analysis, because all items were rated by an equal number of annotators (i.e. 3).

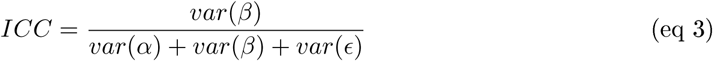

*Kendall’s coefficient* is an association measure based on the ranked items that quantifies the degree of agreement among annotators. As a special case of the correlation coefficient, this coefficient will be high when items’ order (ranked by the given rate) would be similar across annotators. It is based on the computation of the symmetric distances between the ranks, and then normalize it. Because it relies in the distances instead of the absolute values, it better handles consistent rater biases, i.e. the bias effect. In *n*_*c*_ refers to the number of concordant and *n*_*d*_ to the number of discordant with in a sample of *n* number of items.

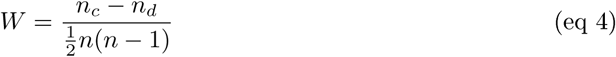

[Gwe02b] demonstrated that Kappa coefficient is influenced by trait prevalence (distribution) and base-rates, thus limiting the comparison across studies. For that reason [Gwe02a] proposed a IRR coefficient (eq 6) that, as Cohen’s Kappa statistic, adjusts the chance agreement raters agree based on a random rating to avoid inflating the agreement probability with not true intentional rater’s agreement. However, *Gwet’s coefficient* has the property of not relying on independence between observations, weights are based on weighted dissimilarities. This coefficient presents several advantages: it is less sensitive to marginal homogeneity and positively biases for trait prevalence (more stable); it can be extended to multiple raters; As Krippendorff’s coefficient it can deal with categorical, ordinal, interval or ratio measures and it can handle missing data; contrary to weighted Kappa, it is not necessary to provide arbitrary weights when applied to ordinal data.

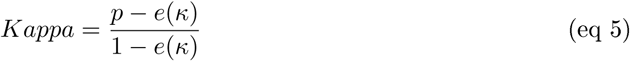

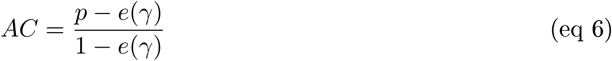

The difference between Gwet and Kappa is in the way that probability of chance agreement are estimated. In Kappa, *e*(*κ*) is based in combining the estimates of the chance that both raters independently classify a subject into category 1 and estimates the probability of independent classification of a subject into category 2 (eq 7). Whereas in Gwet, it is based in the chance that any rater (A or B) classifies an item into a category (eq 8).

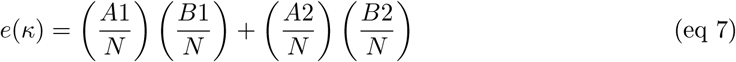

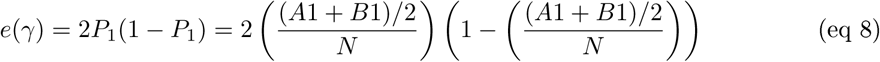

It is important to note that Gwet proposed 2 variants of its statistic, AC1 and AC2. AC2 is a weighted version some disagreements between raters are considered more serious than others of AC1 thus a better alternative for ordinal data. AC2 is intended to be used with any number of raters and an ordered categorical rating system to rate objects, as it is our case. In AC2 both chance-agreement as well as misclassification errors are adjusted, thus it is defined as “bias-adjusted conditional probability that two randomly chosen raters agree given that there is no agreement by chance” ([Gwe02a]).

## 5 Results

### 5.1 Training period

The training period consisted in 4 iterations in each of which a set of sentence pairs was rated by all annotators and afterwards, we had a consensus session where conflicts were resolved and questions about the guidelines were answered, resulting in guidelines updating.

We performed the IRR analysis of each iteration in order to review the effect of consensus sessions in homogenizing the annotation process. As can be seen in figure 3 and table 5.1, the grade of inter-agreement increased in each iteration irrespectively of the statistic. In the 4th session we reached a Fleiss’ Kappa of of 0.546 as the lowest metric, considered as a *moderate* strength of agreement ([LK77]). However, we have to remember that this metric is a non-weighting coefficient, i.e., it considers as bad that an item is rated 0 and 4 by two different annotators, as if they are rated 2 and 3. That is why we reached an *almost perfect* IRR in statistics that better deals with ordinal scales: ICC (0.964) and Gwet’s AC2 (0.910). It is worth to note that Gwet’s coefficients are much more recommended methods to compute IRR than those of the Kappa coefficients family.

**Figure 2:**
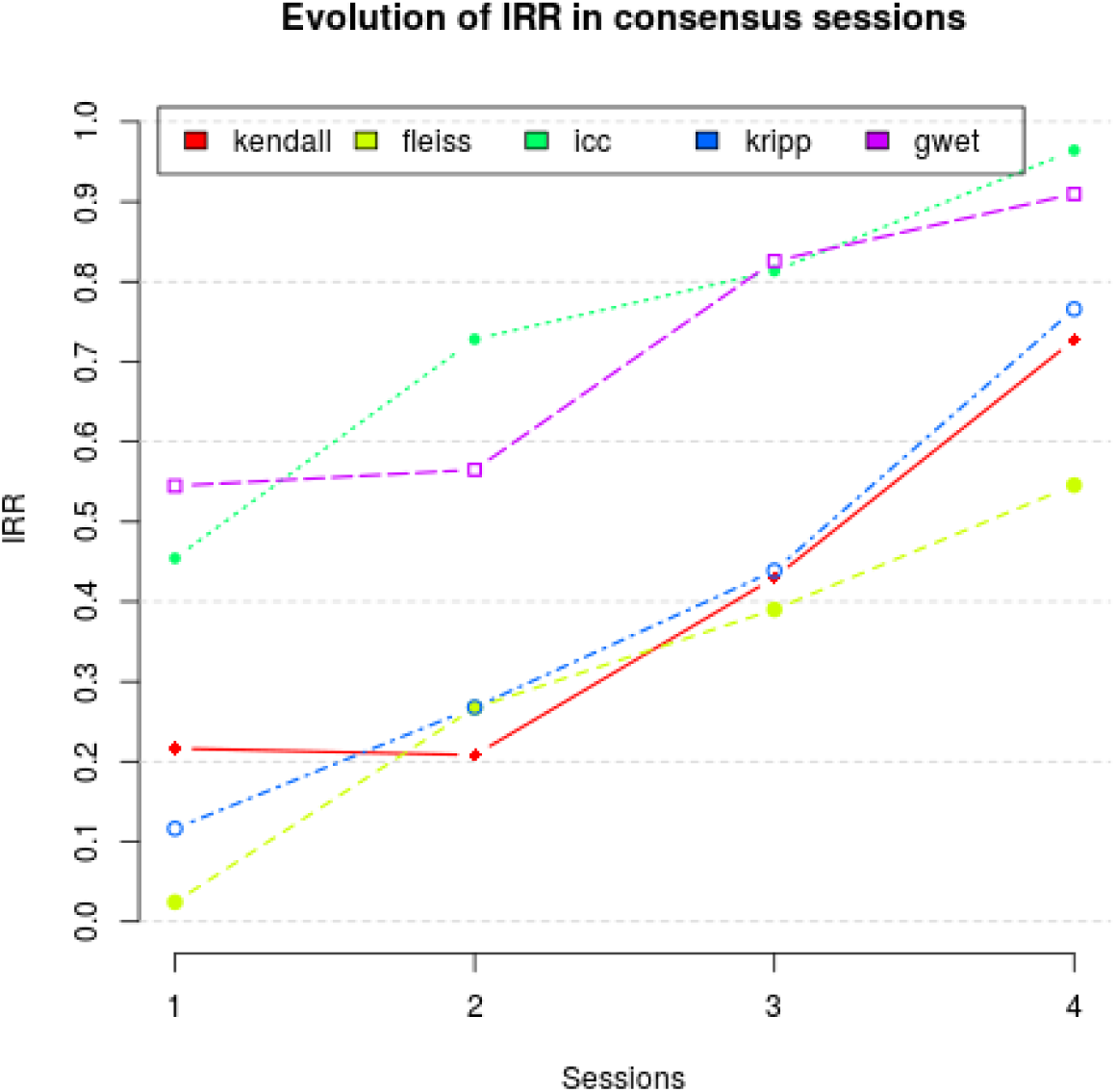
IRR through agreement sessions

**Figure 3:**
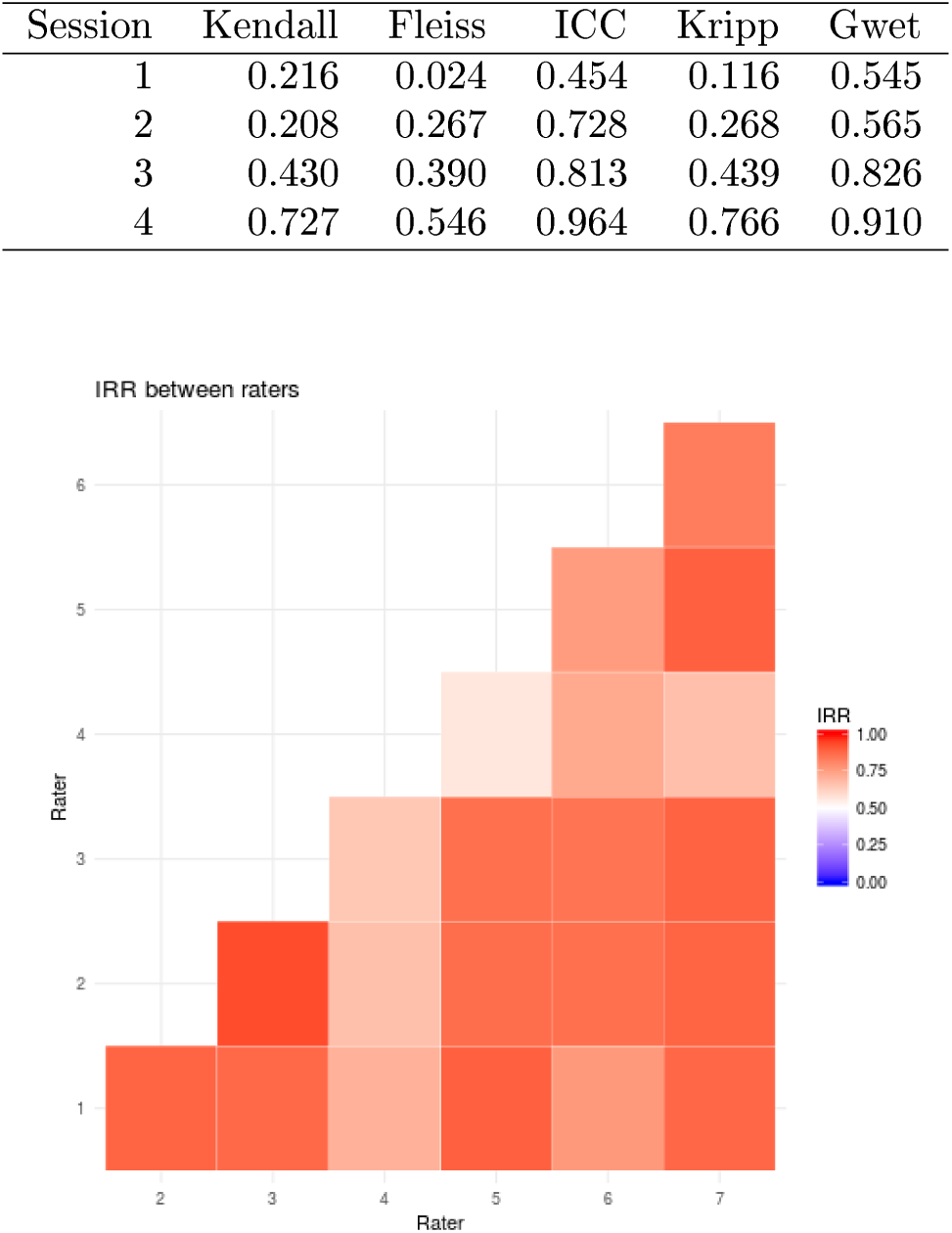
IRR between annotators

We also compared the IRR between all combinations of annotators’s pairs as a way to detect consistent bias of one annotator versus the others, see figure 3. We detected that more guidelines’ clarifications were needed with annotator 4 who consistently had lower IRR with the other raters.

### 5.2 Corpus

After the training period we built the corpus based on the proposed designed (3.2). It resulted in 171 pairs of sentences each rated by 3 annotators taken by chance from a group of 7^7^.

Several IRR analysis were performed to assess the degree that annotators consistently assigned similarity ratings to sentence pairs (see table 2). The marginal distributions of similarity ratings did not indicate a considerable bias among annotators (figure 5) but it shows a prevalence effect towards lower similarity rates (figure 4). A statistic less sensitive to this effect is Gwet’s AC which makes an appropriate index of IRR. in particular the AC2 variant due to the ordinal nature of our data. The resulting coefficient indicated *very good agreement* ([WWWG13]) of *AC*2 = 0.8696 with a 95% confidence interval (0.8399 - 0.8993).

**Table 2:**
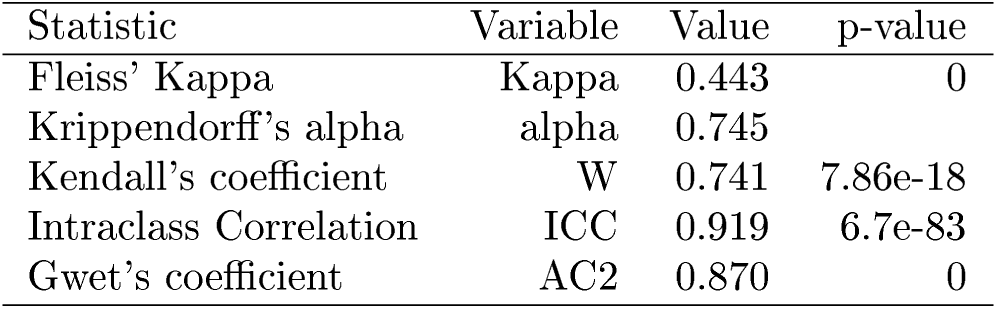
Corpus’ inter-rate agreement for various statistics

| Statistic | Variable | Value | p-value |
| --- | --- | --- | --- |
| Fleiss’ Kappa | Kappa | 0.443 | 0 |
| Krippendorff’s alpha | alpha | 0.745 |  |
| Kendall’s coefficient | W | 0.741 | 7.86e-18 |
| Intraclass Correlation | ICC | 0.919 | 6.7e-83 |
| Gwet’s coefficient | AC2 | 0.870 | 0 |

**Figure 4:**
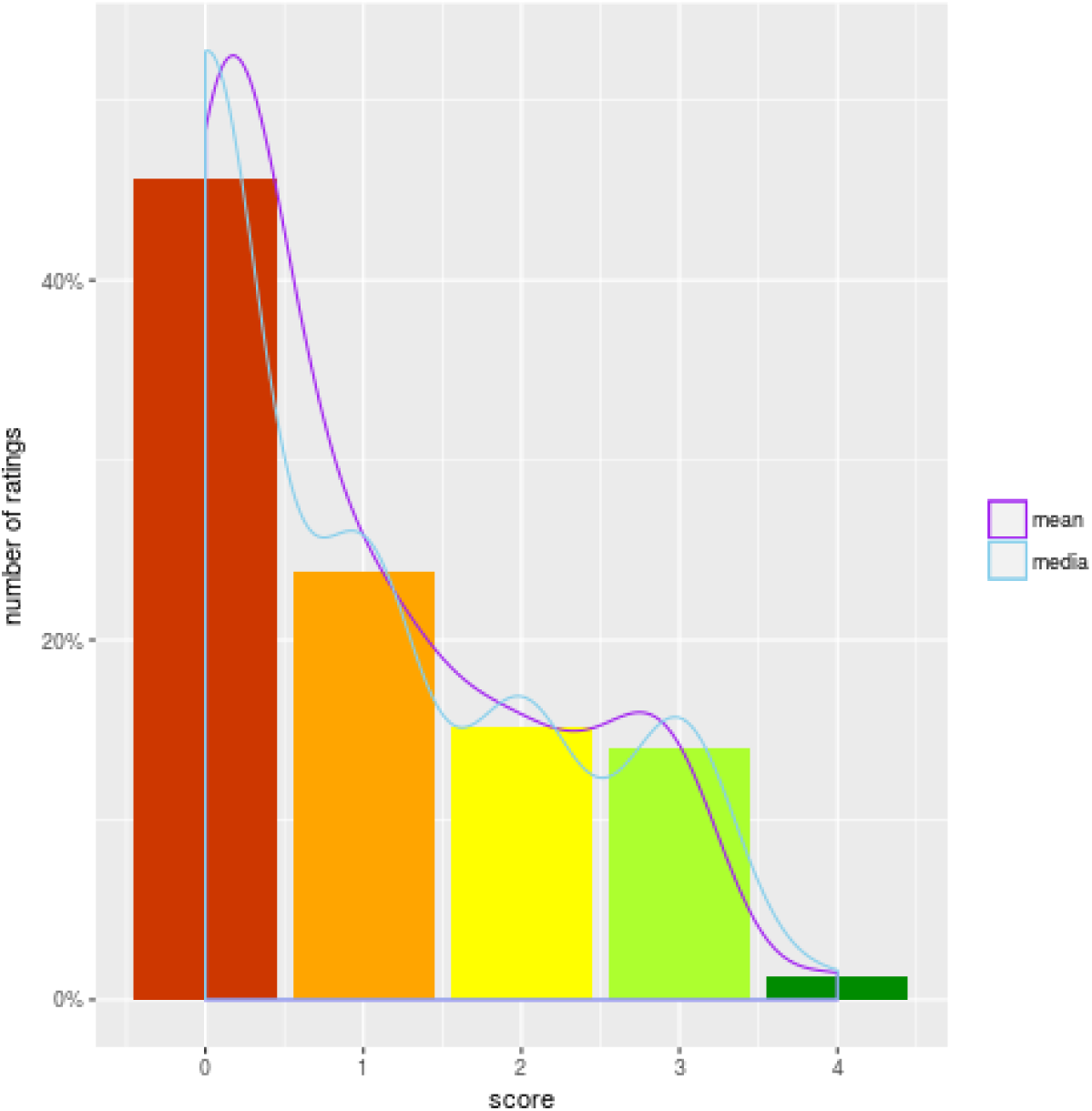
Ratings distribution

**Figure 5:**
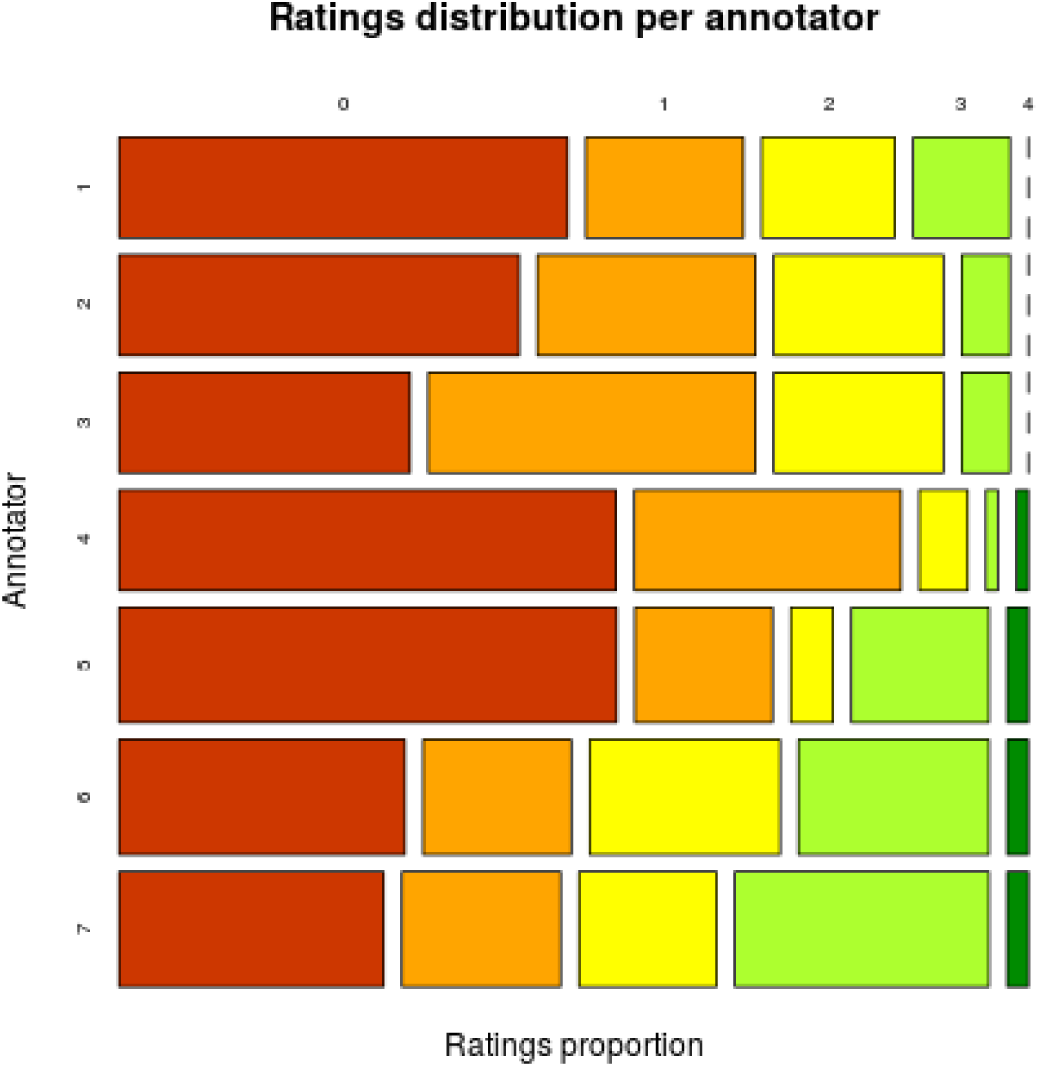
Ratings distribution por annotator

For the sake of clarity, we investigated if the non fully-crossed design caused too inflated coefficients. To do this we first grouped the sentence pairs by the annotators who rate them; next, we computed the IRR for each group it is worth to note that each of these groups is now fully-crossed ; and finally, we computed the arithmetic mean of all groups. The resulting averages (table 5.2) were quite similar to coefficients computed for the whole corpus, re-confirming the corpus reliability.

From the individual ratings distribution (fig 4) we can see that although the distribution is biased towards no similarity, we got a good amount (> 50%) of sentence pairs rated within 1-3 score.

**Table 3:**
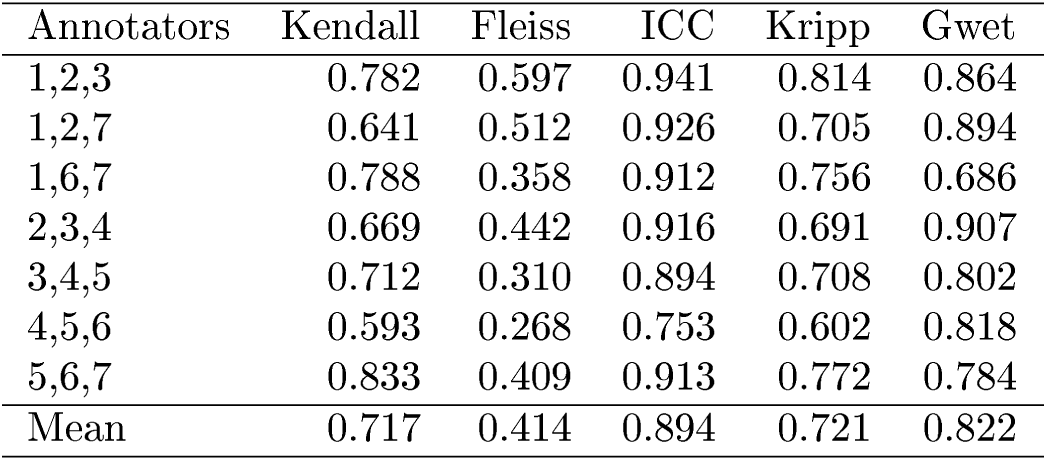
IRR by annotators group

| Annotators | Kendall | Fleiss | ICC | Kripp | Gwet |
| --- | --- | --- | --- | --- | --- |
| 1,2,3 | 0.782 | 0.597 | 0.941 | 0.814 | 0.864 |
| 1,2,7 | 0.641 | 0.512 | 0.926 | 0.705 | 0.894 |
| 1,6,7 | 0.788 | 0.358 | 0.912 | 0.756 | 0.686 |
| 2,3,4 | 0.669 | 0.442 | 0.916 | 0.691 | 0.907 |
| 3,4,5 | 0.712 | 0.310 | 0.894 | 0.708 | 0.802 |
| 4,5,6 | 0.593 | 0.268 | 0.753 | 0.602 | 0.818 |
| 5,6,7 | 0.833 | 0.409 | 0.913 | 0.772 | 0.784 |
| Mean | 0.717 | 0.414 | 0.894 | 0.721 | 0.822 |

## 6 Conclusion

We did not got a corpus with as balanced ratings as desired, however, we have a good representation of 4 of the 5 rates and a corpus with very good inter-rater reliability. Therefore, it is going to serve well our purposes and we think it can be a quite valuable starting point, with respect to data and processes to continue building a standard similarity corpus in the transcriptional regulation’s literature field. To the best of our understanding, this is the first similarity corpus in the topic and thus, it represents a step stone towards the evaluation and training of NLP-based high-throughput curation’s systems.

## 7 Acknowledgments

We acknowledge funding from UNAM, from FOINS CONACyT Fronteras de la Ciencia [project 15], and from the National Institutes of Health (grant number 5R01GM110597-03). CMA is a doctoral student from Programa de Doctorado en Ciencias Biomédicas, Universidad Nacional Autónoma de México (UNAM) and received fellowship 576333 from CONACYT.

## A APPENDIX

### A.1 Contingency tables

**Table 4:** Evaluator-Scores contingency table

|  | 0 | 1 | 2 | 3 | 4 |
| --- | --- | --- | --- | --- | --- |
| 1 | 37 | 13 | 11 | 8 | 0 |
| 2 | 33 | 18 | 14 | 4 | 0 |
| 3 | 24 | 27 | 14 | 4 | 0 |
| 4 | 41 | 22 | 4 | 1 | 1 |
| 5 | 47 | 13 | 4 | 13 | 2 |
| 6 | 27 | 14 | 18 | 18 | 2 |
| 7 | 25 | 15 | 13 | 24 | 2 |
| Sum | 234 | 122 | 78 | 72 | 7 |

**Table 5:** Evaluator’s score distribution

|  | 0 | 1 | 2 | 3 | 4 |
| --- | --- | --- | --- | --- | --- |
| 1 | 0.54 | 0.19 | 0.16 | 0.12 | 0.00 |
| 2 | 0.48 | 0.26 | 0.20 | 0.06 | 0.00 |
| 3 | 0.35 | 0.39 | 0.20 | 0.06 | 0.00 |
| 4 | 0.59 | 0.32 | 0.06 | 0.01 | 0.01 |
| 5 | 0.59 | 0.16 | 0.05 | 0.16 | 0.03 |
| 6 | 0.34 | 0.18 | 0.23 | 0.23 | 0.03 |
| 7 | 0.32 | 0.19 | 0.16 | 0.30 | 0.03 |

**Table 6:** Distribution of scores across evaluators

|  | 0 | 1 | 2 | 3 | 4 |
| --- | --- | --- | --- | --- | --- |
| 1 | 0.16 | 0.11 | 0.14 | 0.11 | 0.00 |
| 2 | 0.14 | 0.15 | 0.18 | 0.06 | 0.00 |
| 3 | 0.10 | 0.22 | 0.18 | 0.06 | 0.00 |
| 4 | 0.18 | 0.18 | 0.05 | 0.01 | 0.14 |
| 5 | 0.20 | 0.11 | 0.05 | 0.18 | 0.29 |
| 6 | 0.12 | 0.11 | 0.23 | 0.25 | 0.29 |
| 7 | 0.11 | 0.12 | 0.17 | 0.33 | 0.29 |

### A.2 Corpus

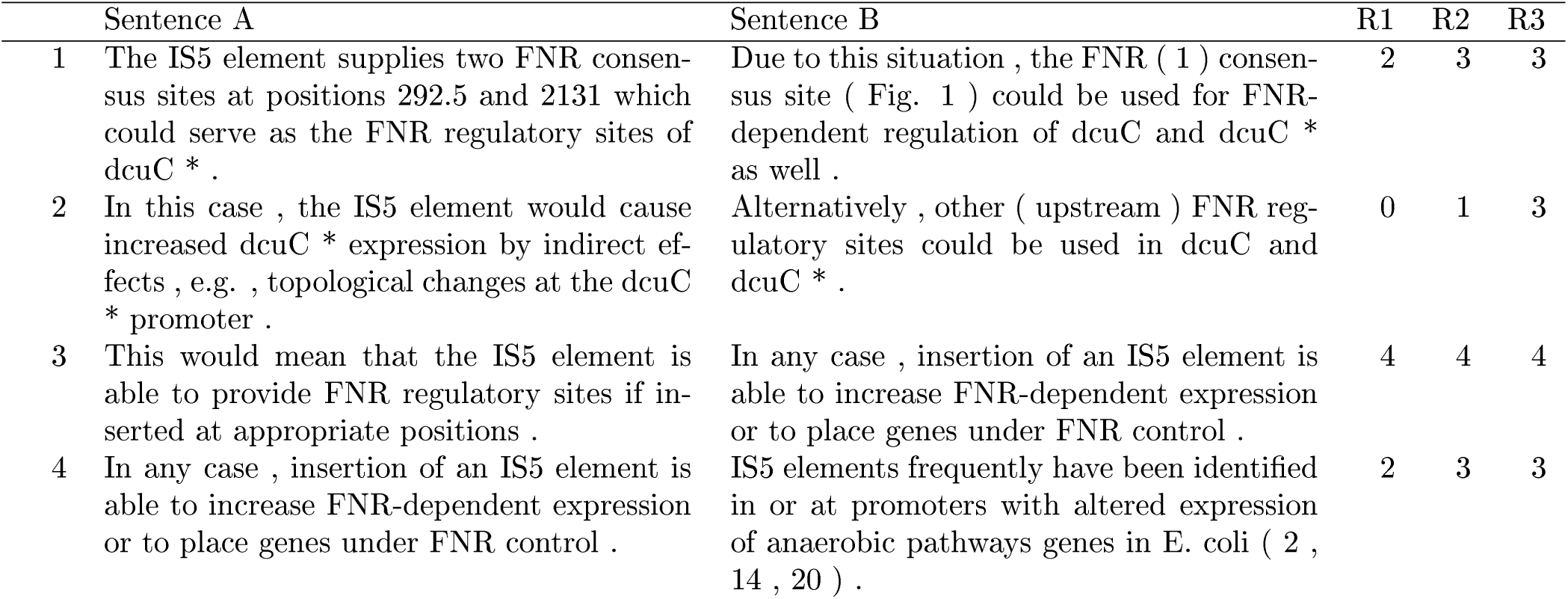
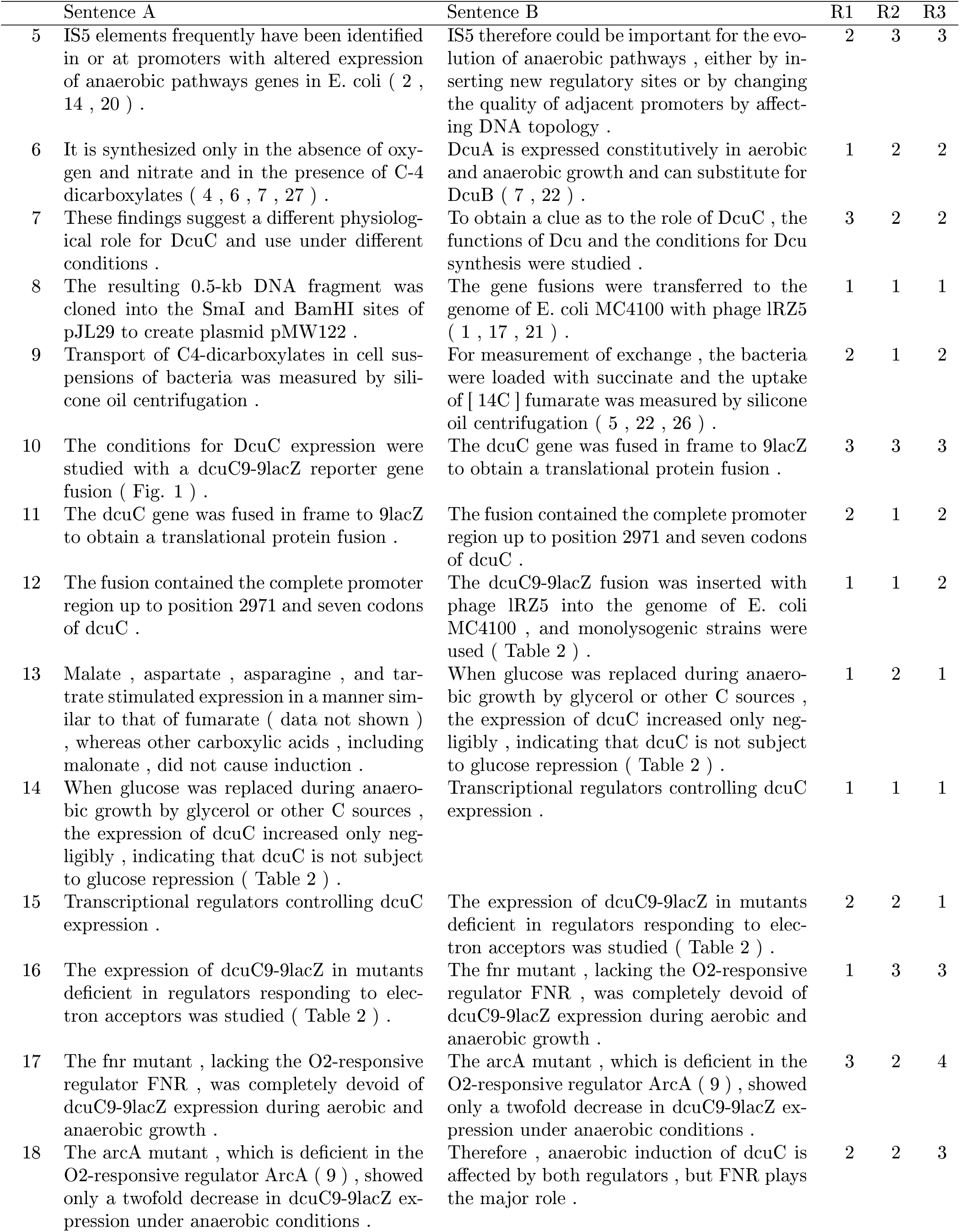
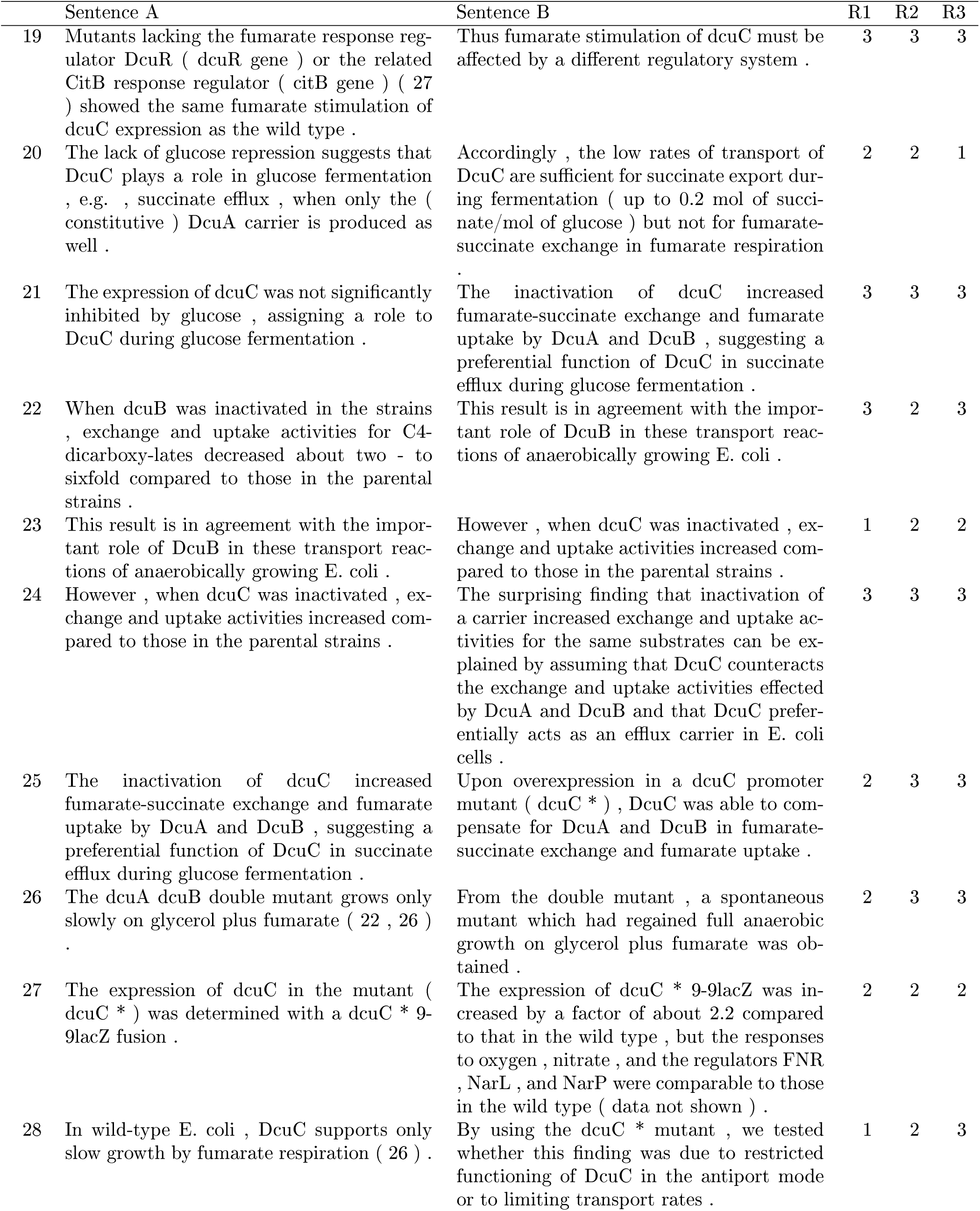
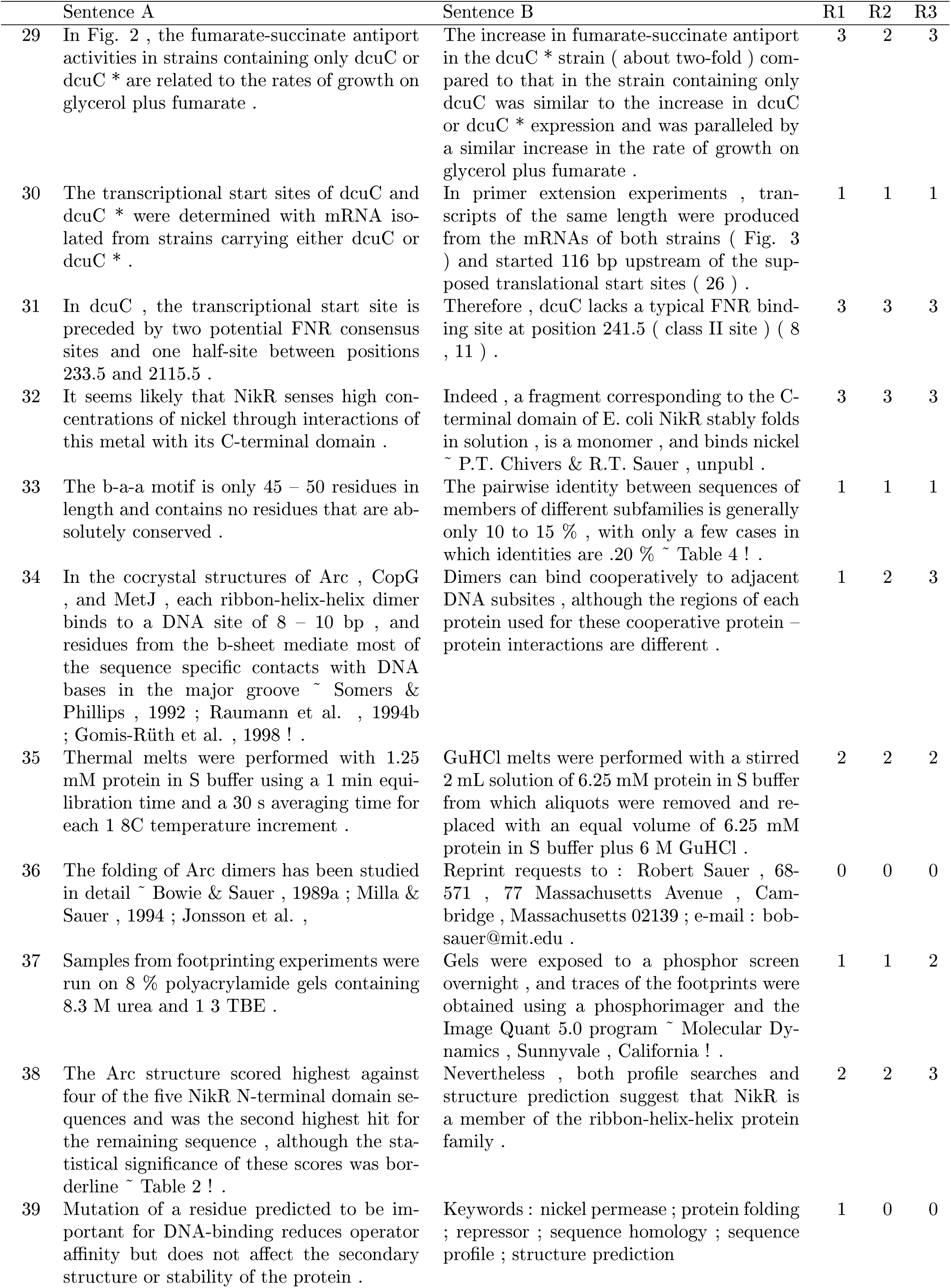
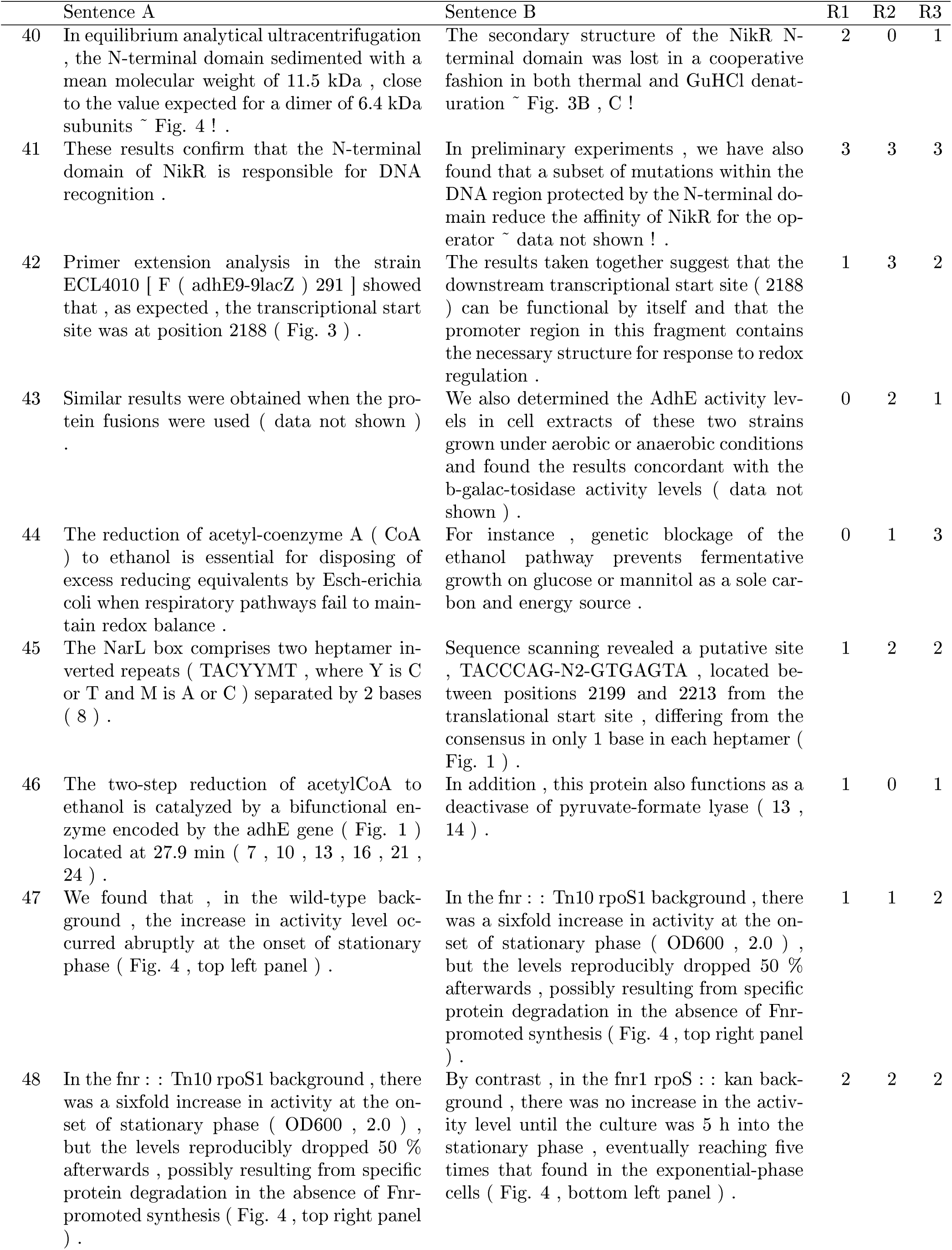
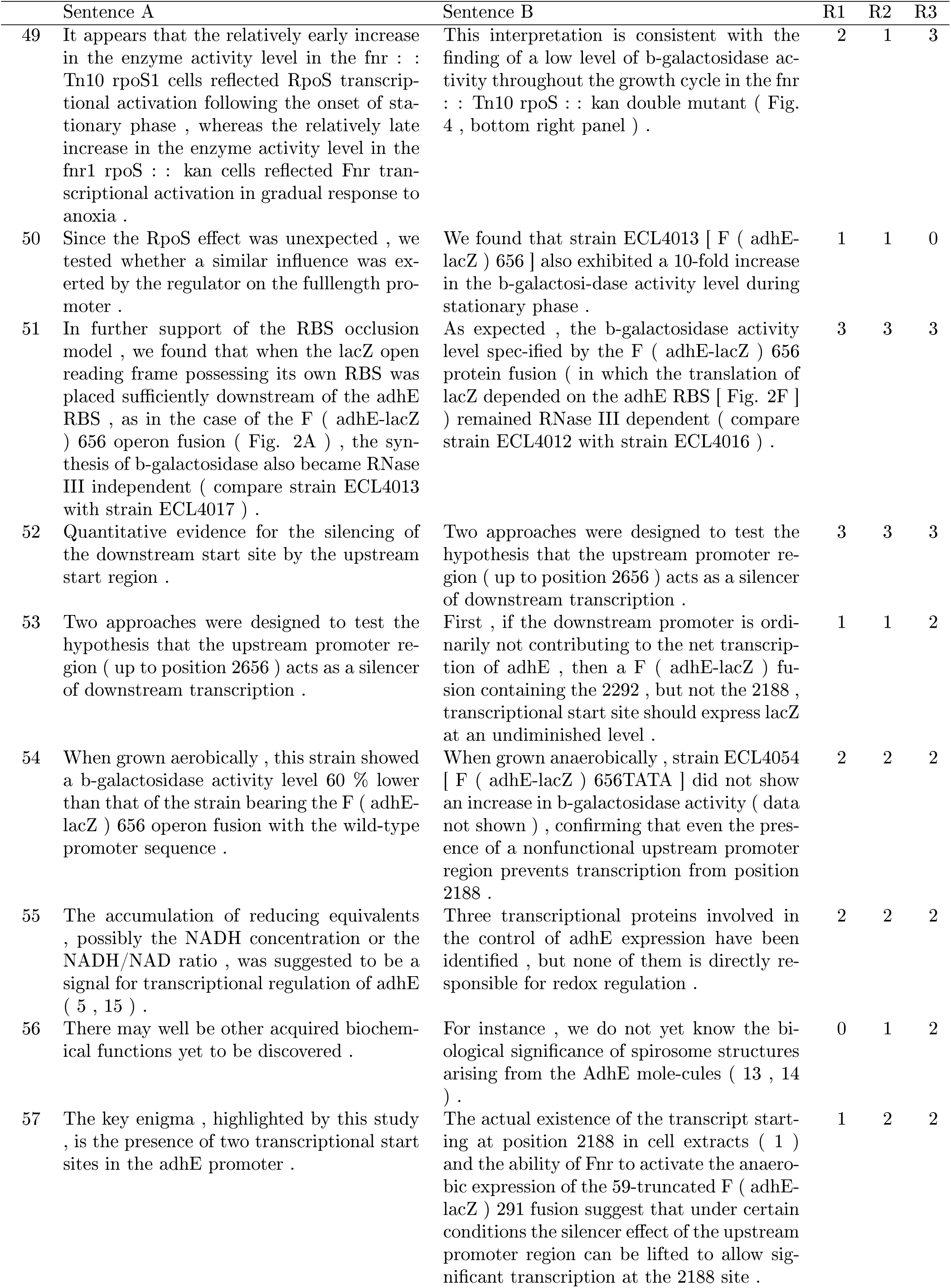
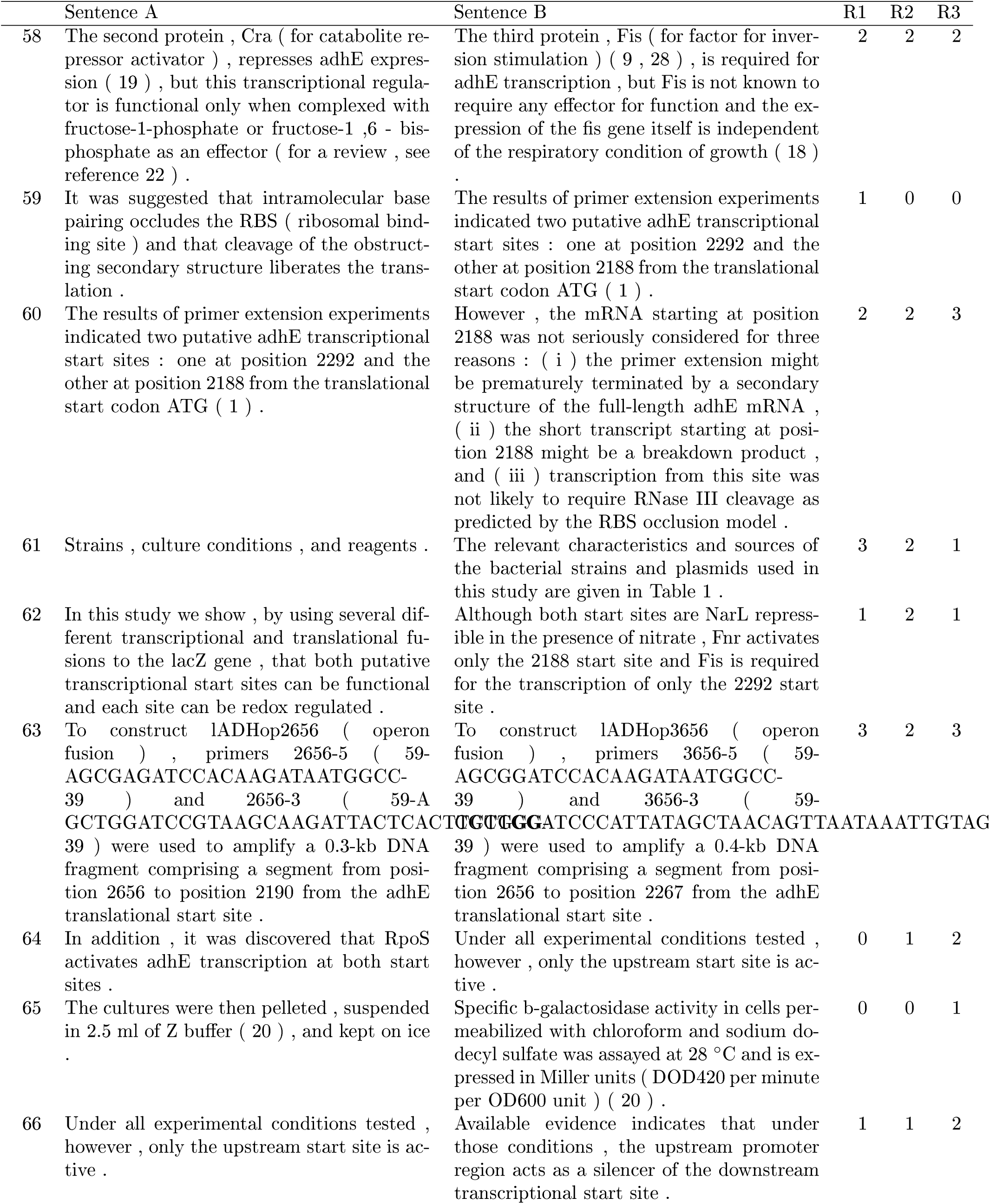
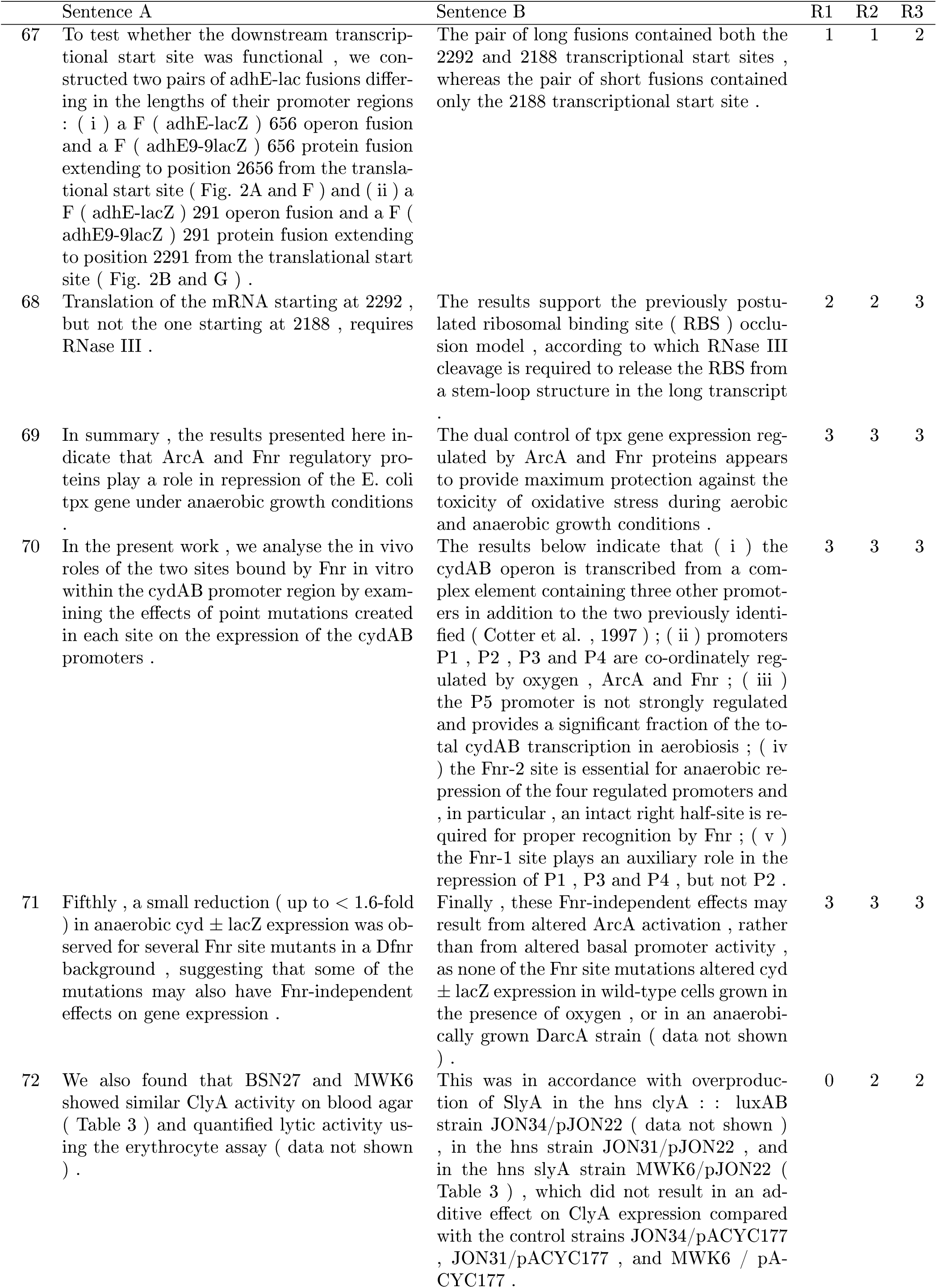
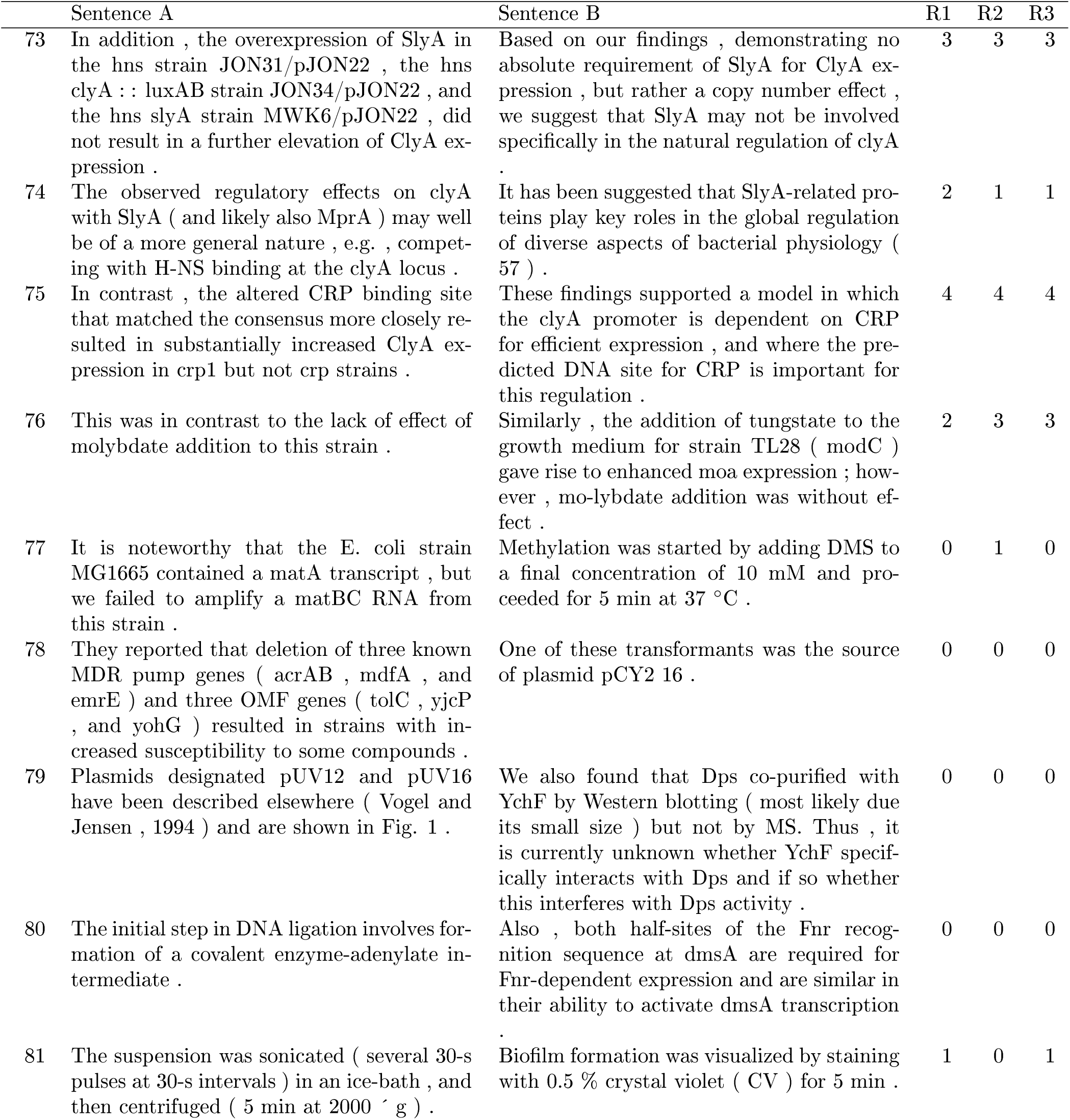
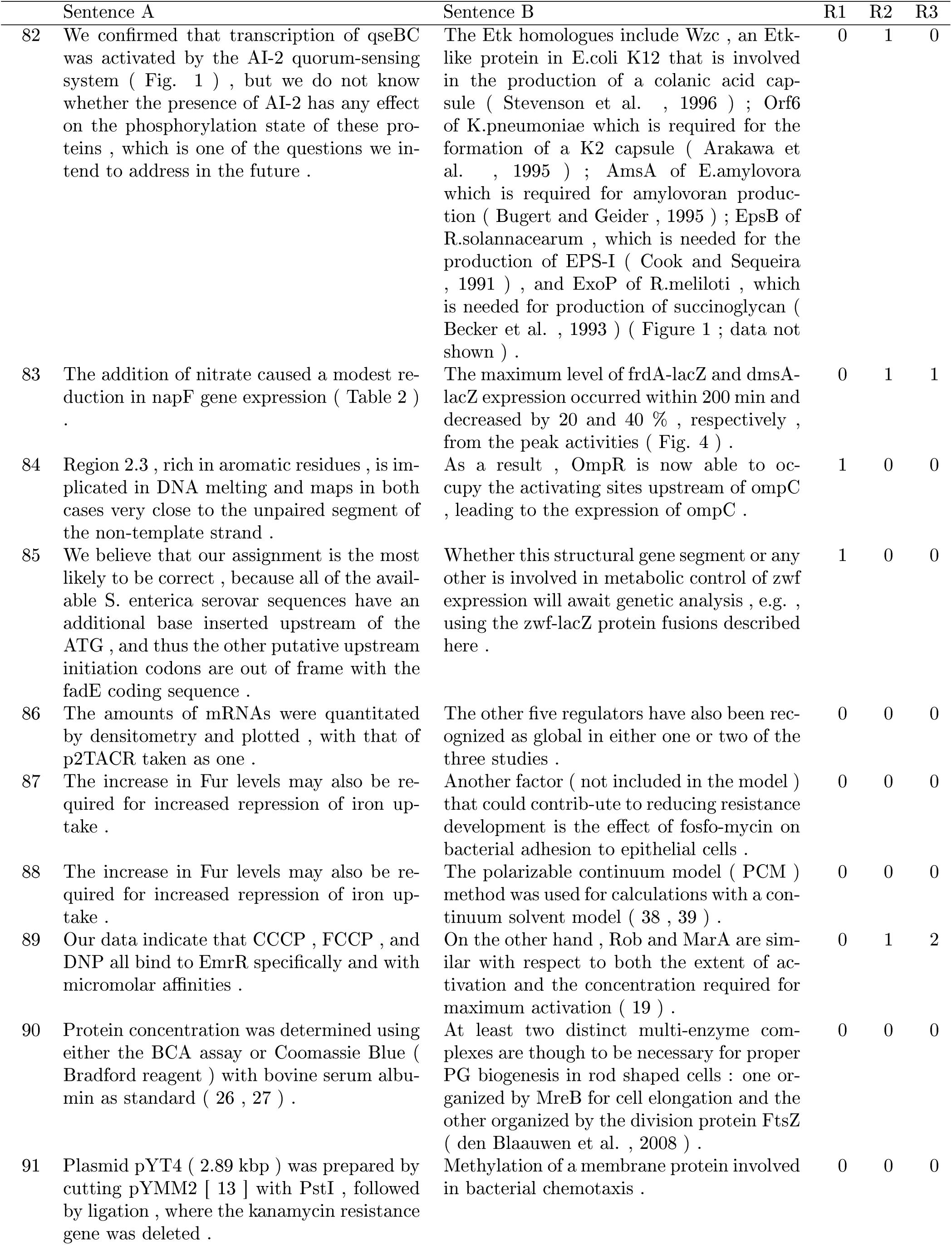
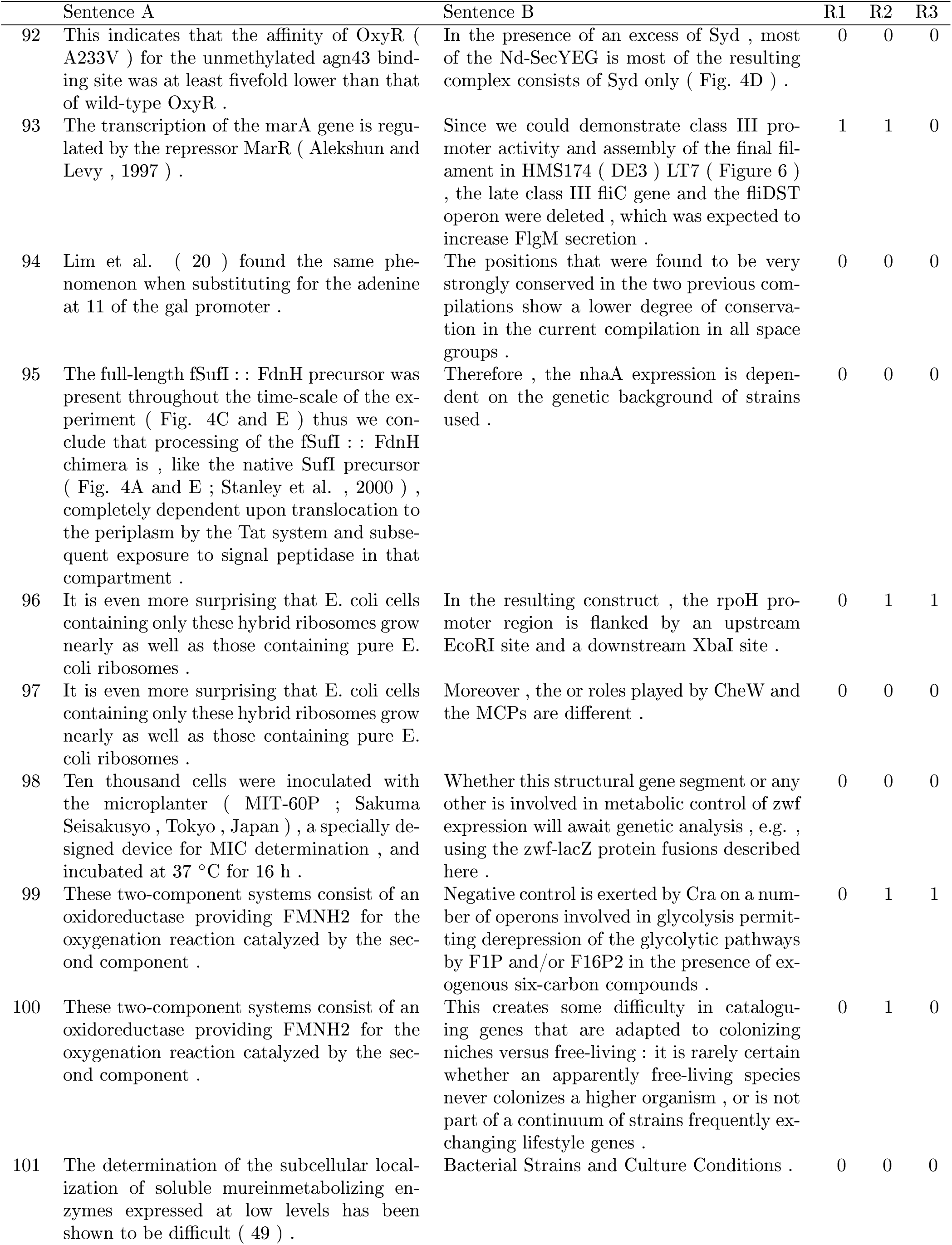
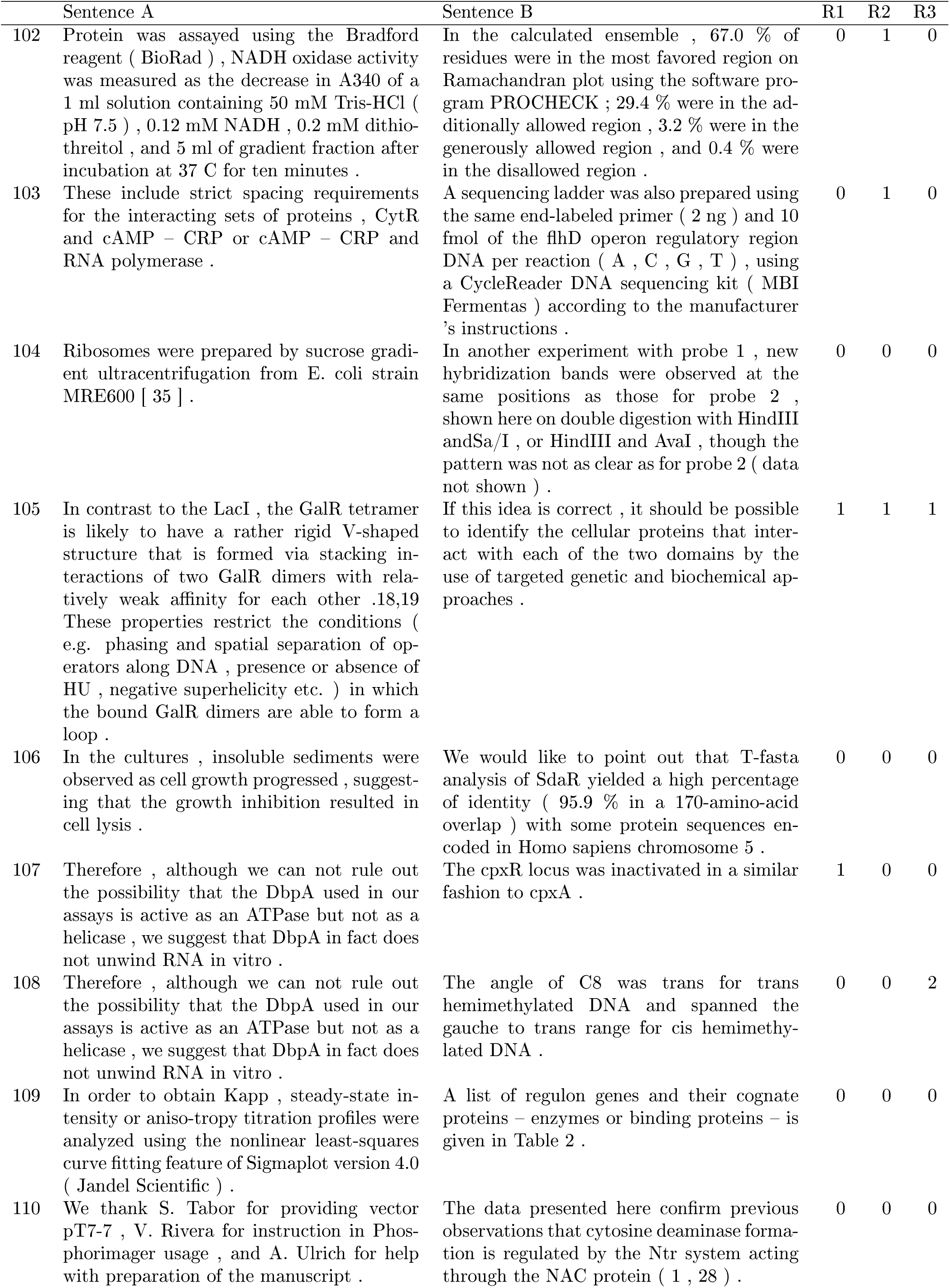
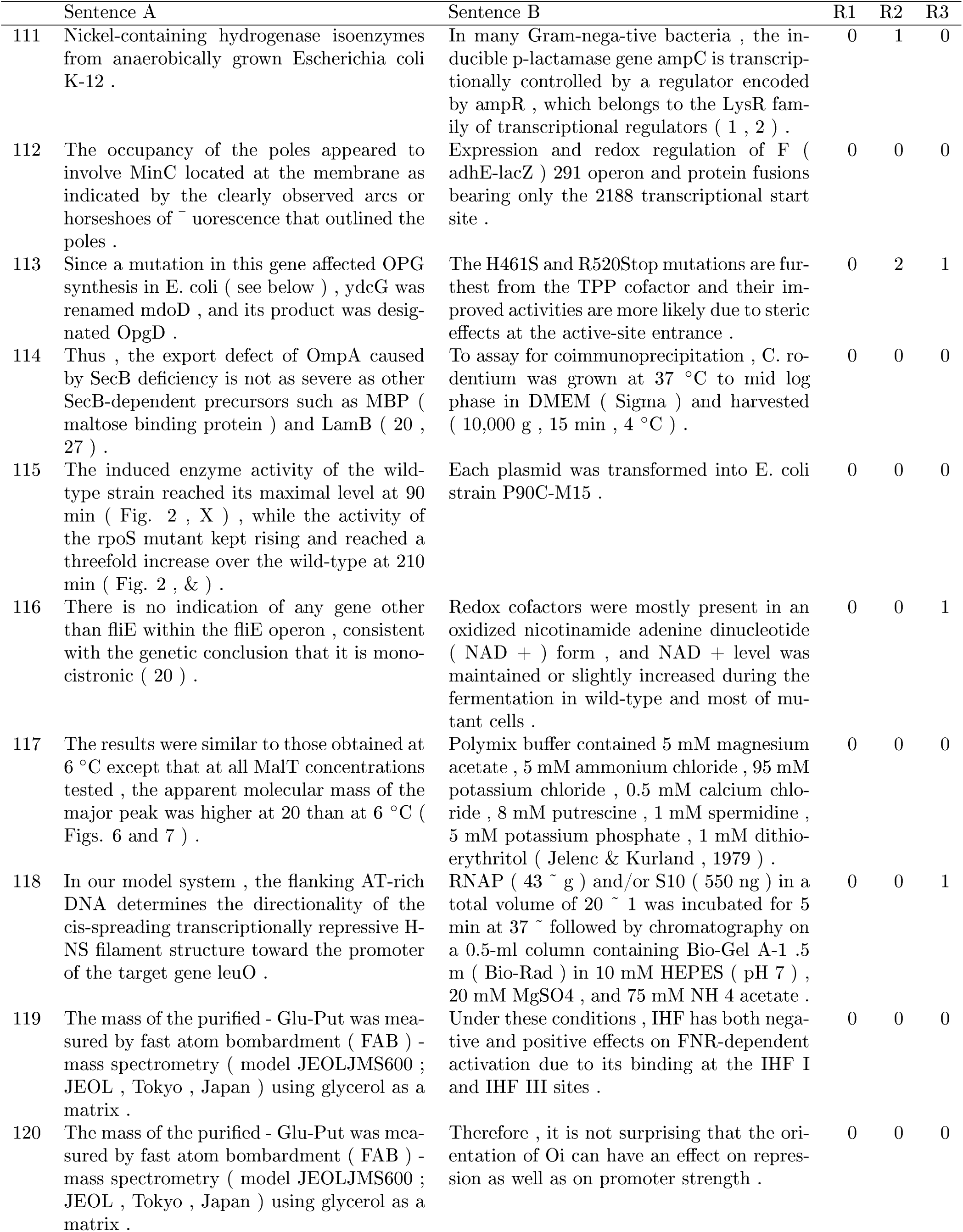
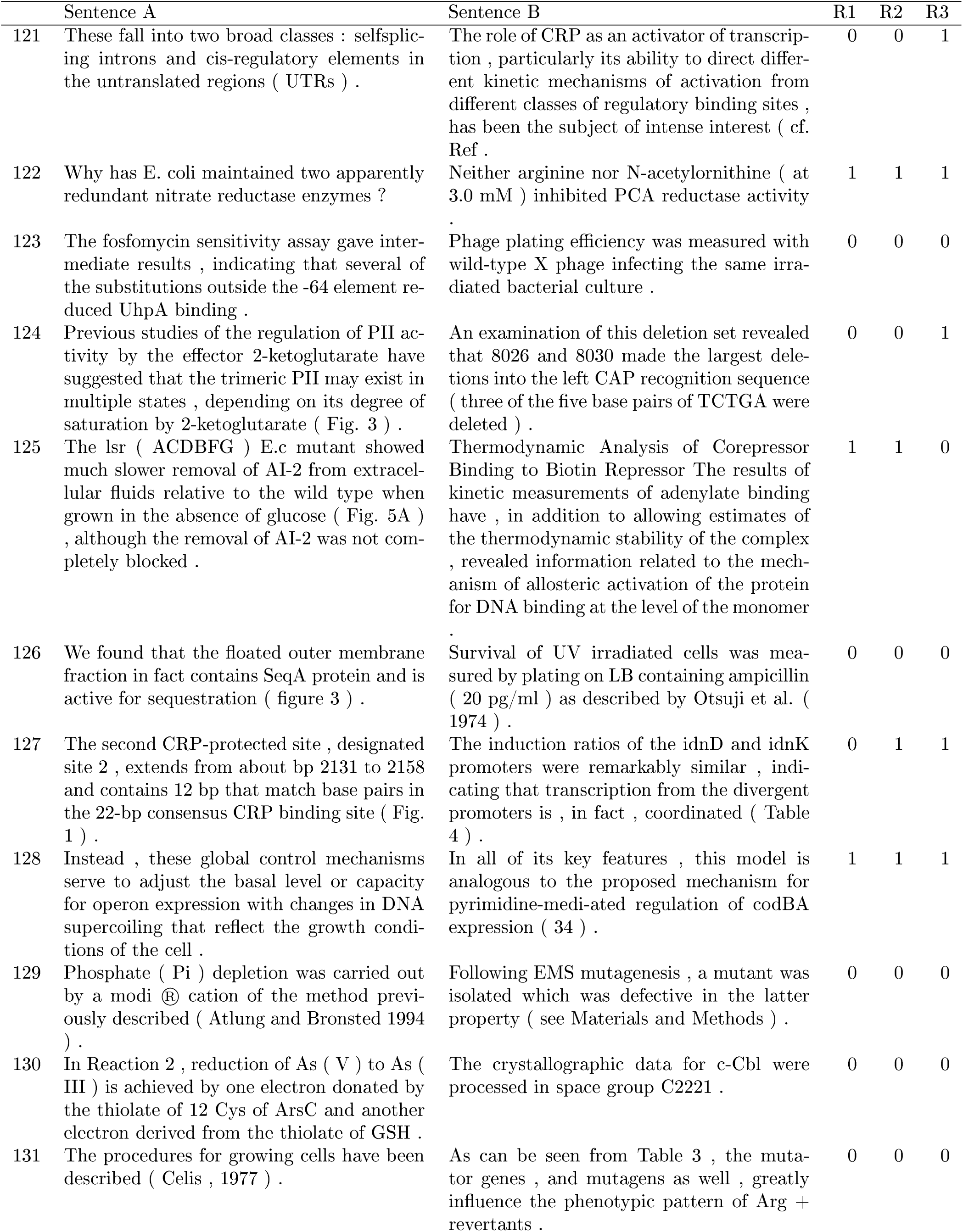
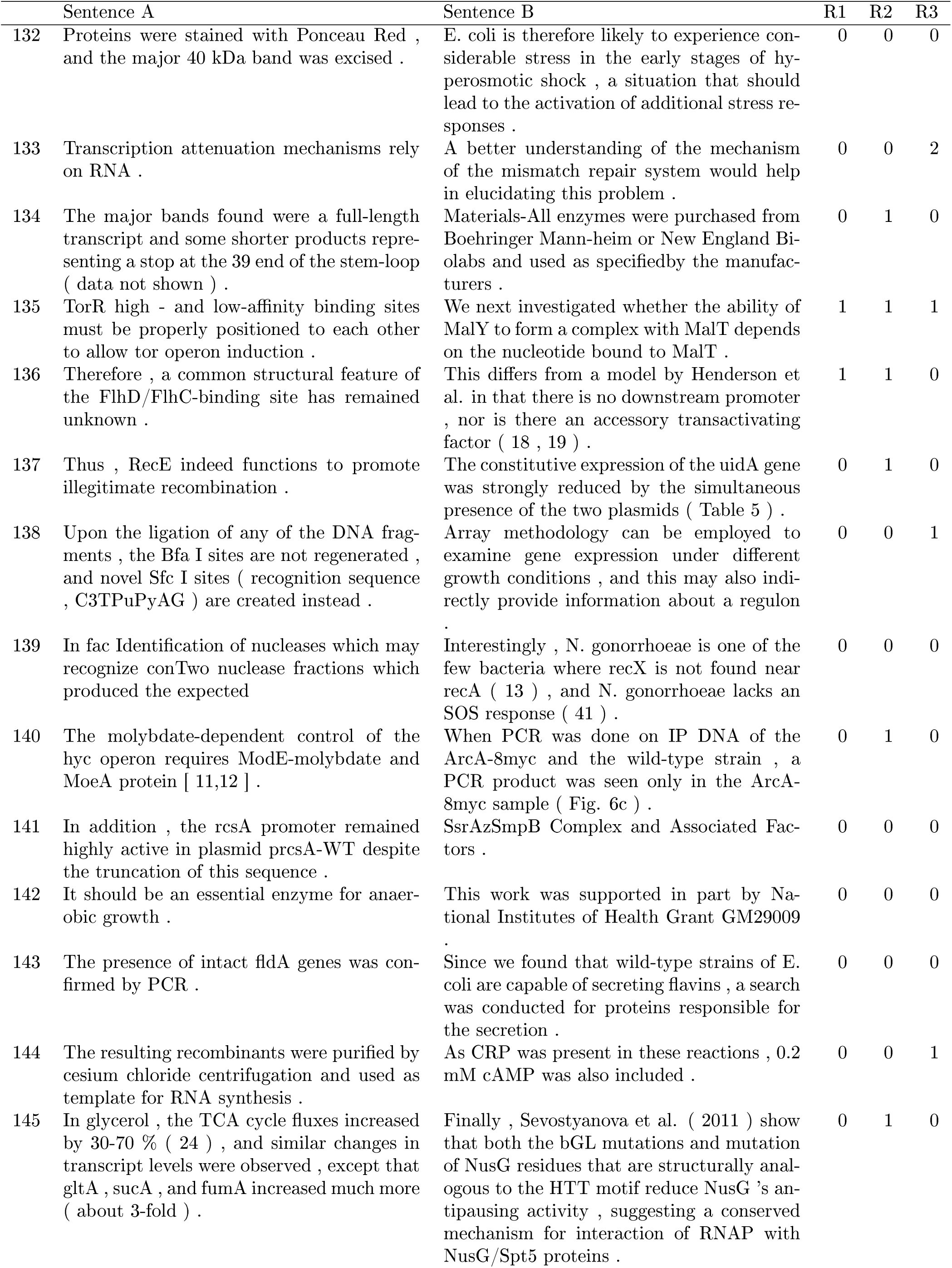
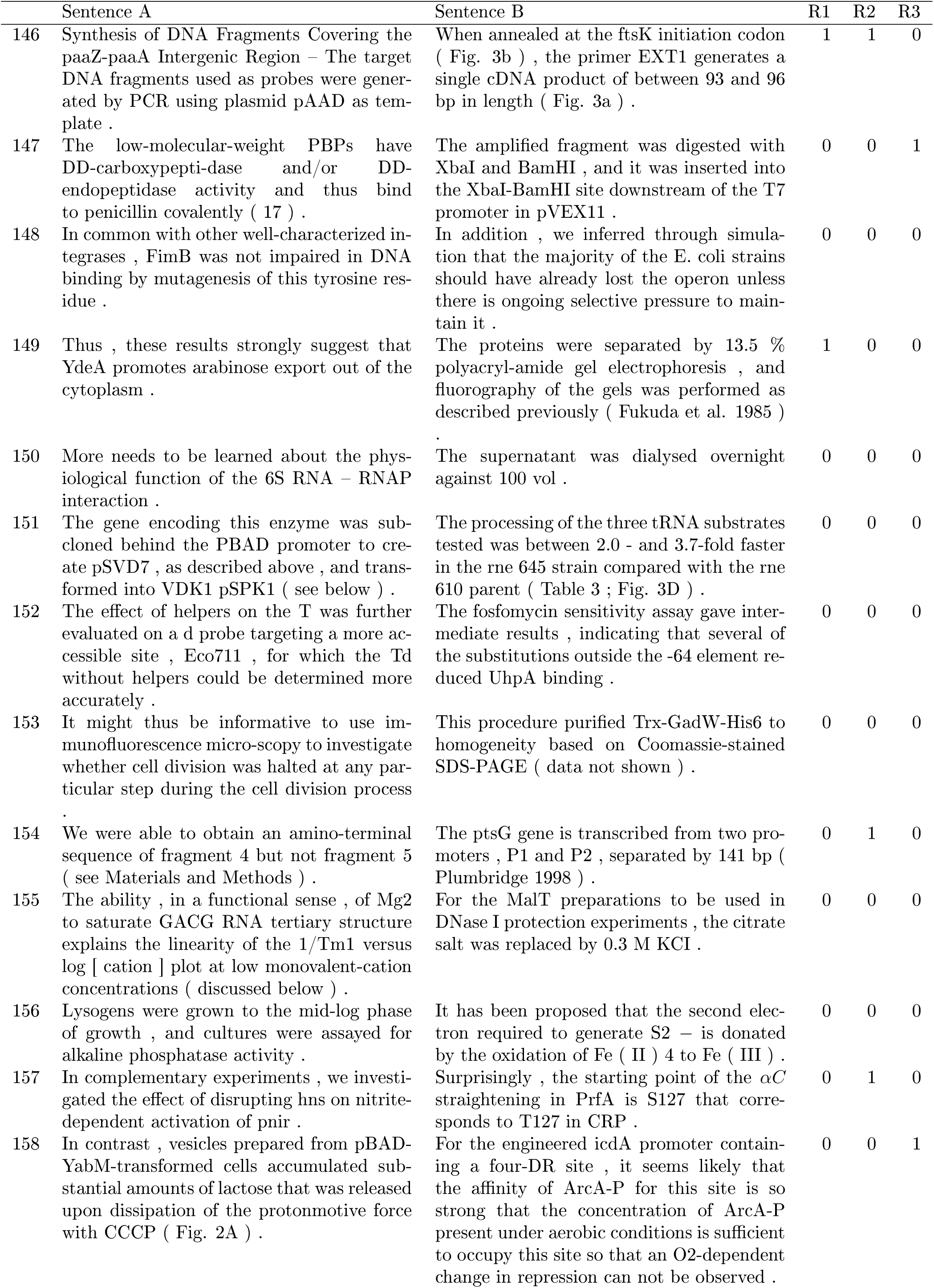
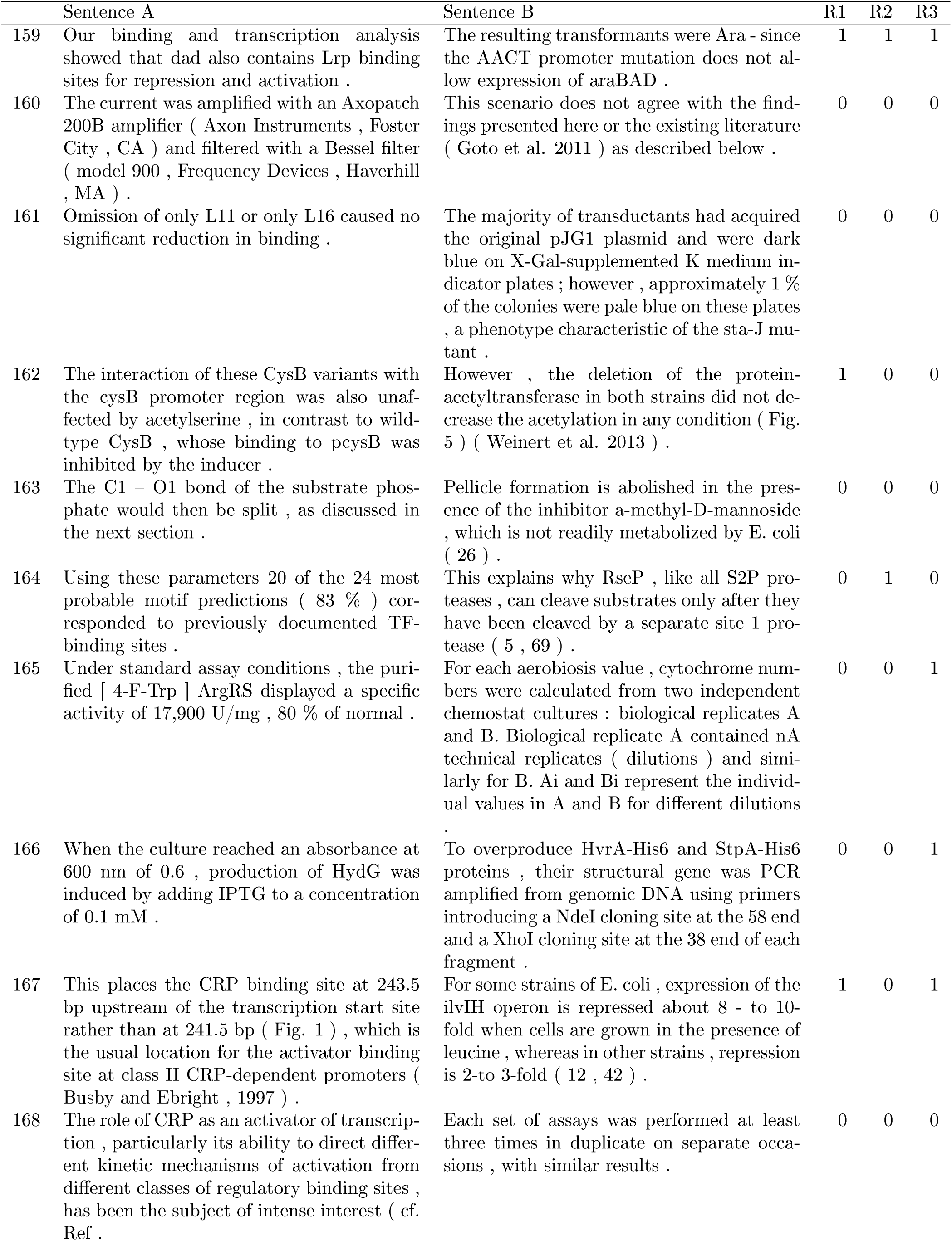
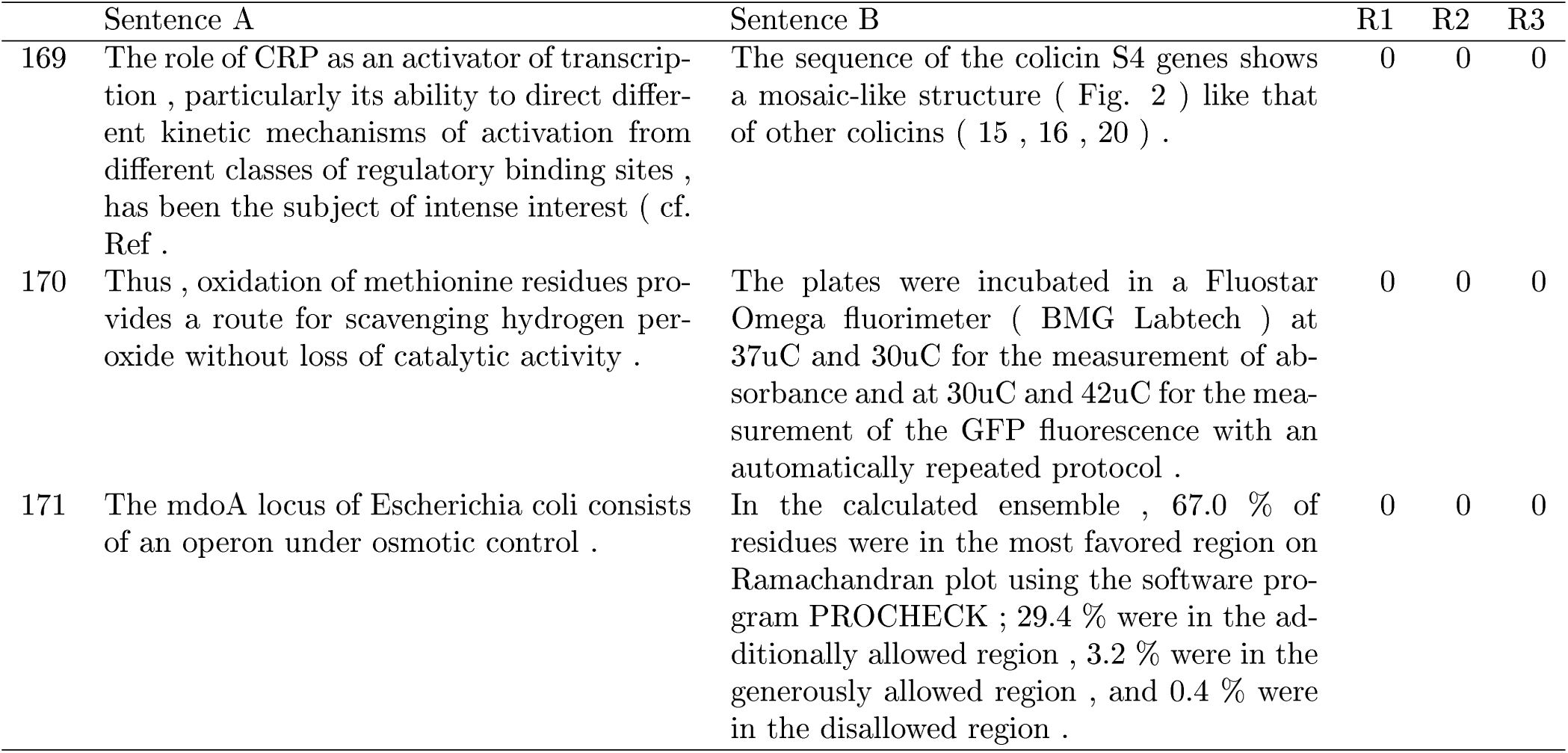

## Footnotes

1 http://rogulondb.ccg.unam.mx/

2 declarative, interrogative, exclamatory, etc.

3 Based on the stvlographic tag assigned by our homo-made PDF processing tool.

4 Applying a baseline metric

5 This is. wo discarded as selection candidates the first 30% and the last 30% of the article’s sentences

6 The 7 groups were taken by chance from all group combinations.

7 Corpus is also available at https://github.eom/JCollado-NLP/Corpus-Transcriptional-Regulation

